# Microbiome Integrity Protects Against Glial-Mediated Tau and Amyloid Pathology Through Circadian and Autophagy Homeostasis

**DOI:** 10.64898/2026.05.20.726549

**Authors:** Kishore Madamanchi, Srinath Gurrala, John Watson, Girish Melkani

## Abstract

Alzheimer’s disease (AD) is characterized not only by tau and amyloid-β aggregation but also by systemic disruptions in circadian rhythms, metabolism, and gut–brain communication that exacerbate neuroinflammation and neurodegeneration. While glial cells play central roles in inflammatory signaling and proteostasis, the contribution of the gut microbiome to glia-driven AD pathology remains poorly understood. Here, we used Drosophila models with glial-specific expressions of human tau and amyloid-associated transgenes to investigate how microbiome integrity influences disease progression. AD models exhibited significant shifts in gut microbial composition, particularly in Lactobacillus and Acetobacter species, suggesting an adaptive microbial response to pathological stress. Strikingly, microbiome depletion (axenic condition) markedly worsened behavioral and physiological outcomes, including disrupted sleep–circadian rhythms, impaired memory, and reduced locomotor function. These deficits were accompanied by amplified neuroinflammatory signaling (Upd–Dome–Hop–Stat92e axis), increased apoptotic gene expression, lipid dysregulation, and altered synaptic markers. Moreover, microbiome loss induced energy stress marked by elevated phospho-AMPK (p-AMPK), yet failed to restore proteostasis, as evidenced by accumulation of ubiquitinated proteins and the autophagy adaptor Ref2p, indicating impaired autophagic flux. This dysfunction correlated with increased tau, phospho-tau, and Aβ42 accumulation. Together, our findings demonstrate that microbiome depletion exacerbates glial-mediated inflammation, disrupts circadian and metabolic homeostasis, impairs, and accelerates cognitive and motor decline. This work highlights a previously underappreciated role of the gut microbiome in restraining glial dysfunction and mitigating AD-like pathology, positioning microbial homeostasis as a critical modulator of neurodegenerative disease progression.

## 1. Introduction

Alzheimer’s disease (AD) is a progressive neurological disorder, and a leading cause of dementia in elderly^1^. AD affects nearly 55 million people worldwide^2^. While the exact cause of AD is not known, several factors contribute to disease severity including genetic, lifestyle, aging and environmental factors significantly contributing to protein aggregation in the brain^3^. Protein aggregates formed by accumulation of amyloid beta 42 (a*β*42) cleaved by *β*-secretase and *γ*-secretase activity on amyloid precursor protein (APP) through amyloidogenic pathway^4^. In addition to a*β*42, microtubule associated protein tau (MAPT) get hyperphosphorylation and forms a tau tangle known as neurofibrillary tangles (NFT)^5,6^. The accumulation of tau and a*β*42 contribute to neuronal loss and brain shrinkage^7^. Protein aggregation significantly impacts the memory, and behavioral activities^8,9^. Despite significant research focusing on understanding etiology of AD, an effective treatment paradigm to slow down or prevent progression is still unclear. In addition to a*β*42 and NFT accumulation, in patient’s chronic neuroinflammation acts as an important factor in driving AD pathology^10^. Activated glial cells such as microglia and astrocytes release proinflammatory cytokines that propagate neuronal stress and synaptic dysfunction^11^. Increased neuroinflammation reduced anti-inflammatory responses further reinforce the disease progression^12^. These molecular disruptions further contribute to physiological behavioral changes^13^, such as alterations in sleep-wake cycles^14^, memory processing^15^, changes in eating patterns, gut microbiomes in patients^16^. We hypothesize that gut microbiome abundance is key regulator of behavioral and pathophysiological responses, and its absence could worsen AD and associated dementia pathology. Several studies reported the altered microbiome composition significantly impact inflammatory responses by releasing bacterial secondary metabolites and cellular components^17–20^. That significantly influencing AD pathology^21^ by activating the brain resident immune cells and reducing the clearance of aggregated proteins in AD patients^22–24^. Despite growing evidence linking the gut microbiome to AD, the causal mechanisms by which microbiota influence disease progression remain poorly defined. Earlier studies have largely focused on systemic inflammation or neuronal outcomes^25,26^, with limited attention to how gut-derived inflammatory signals directly affect glial-driven amyloid and tau pathology through pathways that integrate circadian rhythms, metabolism, and autophagy. Although lipid dysregulation, sleep disturbances, and impaired proteostasis are now established hallmarks of AD, it remains unclear whether these features arise independently or from a coordinated failure of microbiome-dependent regulatory networks within glial cells. In addition, prior mammalian studies are limited by a lack of cell-type specificity and challenges in microbiome manipulation, hindering mechanistic interpretation. Therefore, a critical knowledge gap persists in understanding how the gut microbiome modulates glial inflammatory signaling and proteostasis pathways to shape the metabolic, behavioral, and pathological outcomes of AD. To understand the role of glial cells and gut microbiome in altering AD pathology, we performed glial specific expression of humanized tauopathy mutations, amyloid precursor protein, and a*β*42 genes. Which could lead to changes in gut microbiome composition followed by alterations in behavioral and memory functions.

To further understand the role of microbiome roles with AD, we have depleted microbiome (axenic flies) and performed multiple behavioral, cytological and biochemical assessments. Axenic lies were generated by growing dechorionated embryos on antibiotic supplemented standard food and these flies termed as axenic antibiotic (AA) flies, and flies without antibiotic food termed as conventional control (CC) flies. Our study found a significant change in *Lactobacillus* and *Acetobacter* species in tauopathy and amyloidopathy models in CC condition, which are critical for maintaining the healthy gut microbiome in AD flies. In addition, lack of gut microbiome (AA) showed severely compromised memory and sleep quality in AD transgenic flies. We also observed an increased protein aggregation, upregulation of neuroinflammatory like *Upd* (unpaired) 1, 2, and 3, *Eiger* and their downstream targets such as *Dome*, *Hop*, *Stat92e* and *Imd* genes. In addition, cell death promoting genes such as *Hid* and *Reaper* which are involved in caspase mediated cell death were also upregulated in AA flies. Overall, our study supports the significance of gut microbiome through its regulation on inflammatory gene suppression, modulating sleep quality, memory and locomotion behaviors which are significantly altered during AD pathogenesis.

## 2. Materials and Methods

### 2.1. *Drosophila* rearing condition and glial specific expression

All the control and experimental flies were maintained in standard diet with active dry yeast 3 g/L, agar 11 g/L, yellow cornmeal 55 g/L, molasses 72 mL/L, propionic acid 6 mL/L, and 10% nipagin 8 mL/L. Flies were housed at 50% humidity at 25 °C in 12 hours light-12 hours dark (LD) cycles. Fly food was changed every 3-4 days^27–30^. *Drosophila* fly lines obtained from Bloomington *Drosophila* stock center (BDSC), and GAL4/UAS system used for cell, and tissue specific gene expressions. For glial specific expression we used *Glaz*-*GAL4* (BL#23353) drivers. Control lines such as *w^1118^* (BL#5905), throughout the paper (+) refers to *w^1118^*, UAS-GFP (BL#5431), human AD transgenic genes under UAS control, such as UAS-APP (amyloid precursor protein, BL#64384), UAS-a*β*42 (BL#33770), UAS-HTau (BL#51362), UAS-hyperphospho HTau (BL#51365), and UAS-HTau*^R406W^* (obtained from Mel B Feany’s lab, Harvard medical school) were used. These driver lines were crossed with control and over expressed lines to augment AD pathology and to understand their impact on experimental flies. Progeny (F1) of all the experimental flies were collected at eclosion and gender separated and maintained in separate vials 25 flies in each. All the flies were maintained till the experiments were completed.

### 2.2. Depletion of microbiome for preparation of axenic flies

Embryo collecting chambers (Flystuff® *Drosophila* Embryo Collection Cages-Genesee Scientific) were used to collect 24 hours old embryos laid by the parental flies and food was changed every 24 hours to collect fresh embryos. Axenic flies were generated by dechorionizing 24 hours old embryos before larva coming out of embryo using 70% ethanol for 3-5 minutes^31^. Once the chorion released the embryos were washed in sterile water and then transferred to standard fresh food and antibiotic cocktail containing food vials. After eclosion food was changed every 3 days throughout the adult stage (up to 3 weeks)^31^. Previously, we used axenic sterile (standard food changed every 24 hours instead of every 3 days to maintain sterile environment) method as well to make microbiome free flies along with antibiotic food after dechorionizing the embryos, where we observed similar results^31^. The axenic fly food was prepared by supplementing antibiotics such as Ampicillin (Cas No.69-52-3), Kanamycin (Cas No.25389-94-0), Tetracycline (Cas No.60-54-8), Erythromycin (Cas No.114-07-8) to the standard diet during food preparation^31^. Each antibiotic was weighed (50 mg/L), dissolved and added to the food at 60 °C after autoclave and mixed thoroughly based on standardized protocol^31–33^. Flies grown on antibiotic food are termed as axenic antibiotic (AA) and flies grown on standard food termed as conventional control (CC). Antibiotic treatment will eliminate the external microbiome and reduce the internal/gut microbiome quantity significantly^31,33^.

### 2.3. Sleep/circadian activity

*Drosophila* sleep/circadian activity behavior was studied using *Drosophila* activity monitor system (DAM, TriKinetics Inc MA, USA) under 12 hours light-12 hours dark cycles at 25 °C for 5 consecutive days^34,35^. We performed sleep analysis for control (*w^1118^*, GFP) and AD (HTau, HTau*^R406W^*, hyperphospho HTau, a*β*42, APP) gene over expression male progeny flies of *Glaz*-*GAL*4 drivers. 2.5-week-old male flies were housed in a glass tube provided with food. We performed sleep/circadian activity analysis for both conventional control (without antibiotics/flies with natural microbiome-here after referred as-CC) and axenic antibiotics (with antibiotics/without microbiome-here after referred as-AA) flies. We used 16 flies from each genotype with three biological replicates (n=3). The vials were labelled with random numbers to prevent bias during setting up of experiment. We measured Sleep, activity during daytime, nighttime and overall (24 hours) for 5 days. Raw data was used to analyze sleep (immobility of a fly >5 minutes), activity (fly crossing of an infrared beam at the center of DAM), sleep bout (inactiveness of fly minimum of five consecutive minutes) bout length (duration of each sleep period/sleep bout)^36^. We measured the sleep quality by the fraction of sleep bouts and bout length known as sleep fragmentation index which provide the details about individual fly sleep quality. Average sleep/activity behavior of individual fly from each genotype and condition in 24 hours (ZT0-ZT24) was calculated using “One-Click sleep analysis software application” developed in python background in our lab.

### 2.4. Locomotion performance using geotaxis assay

To evaluate the locomotion behavior of our experimental flies through geotaxis assay we have designed and developed a 3D-printed motor mounted automatic device to test the locomotory performance. The device is designed with a tube holder which accommodates 12 vials and a Raspberry Pi camera to record and a light source to clearly capture the fly movements. The camera connected to Raspberry Pi 4B monitor to observe and record the assay while performing the assay. The whole system works based on Python script developed in our lab connects and control both the camera and motor of the device simultaneously. We used 3-4 technical replicates with 11 flies per biological replicate (n=3) with random labelling before performing the experiment. Video was recorded after the initial tapping of device to settle the flies at the bottom of the tube/column. For the recorded video (and any additional video from the same experiment), it is processed by a new python script which takes in the video and a faster RCNN deep learning model was applied to detect the vial tube boundaries in the video. Then computer vision techniques are applied such as background subtraction for individual fly subtraction. The Python script collects locomotion behavior across all biological replicates for each vial but taking the mean position of all the flies within a vial over all the frames^37^. This Python script final outputs relevant plots and csv files to assess the performance of the flies and allow for further statistical analysis through tools like PRISM.

### 2.5. *Drosophila* olfactory aversion test for memory assay

The experimental flies were allowed to explore the T-maze (CelExplorer Labs) provided with two neutral odors (3-octanol and 4-methylcyclohexanol) diluted in mineral oil (1:10 ratio). The flies were grouped into 30-40 per experiment from each genotype per biological replicate (n=3). To prevent the bias, we labelled the vials other than actual genotype labelling, and the assay was performed based on standard protocol^38^. The test consists of three phases (Naive-N, Training-T and testing). Naive includes exposing the flies to two different odors in T-maze containing two odor chambers for 3 minutes 30 seconds to determine the odor preferences of flies. Training round consists of 2-minute 30 seconds shock session (100-110 volts) with an odor based on naive results, followed by a non-shock session with second odor for 2 minute 30 seconds (repeated three times). After the shock sessions the flies were allowed to settle for 5 minutes and then expose to two odors simultaneously for 3 minutes 30 seconds then the chambers were closed to retain the flies in the respective odor chamber to score the flies. As previously described and recently established from our lab, performance index was calculated by counting number of flies choosing shock odor, subtracting with flies choosing non-shock odor and divided with total number of flies^39,40^. Avoidance index calculated by dividing the number of flies chosen non-shock odor with total number of experimental flies then multiplied with one hundred^40^.

### 2.6. Cytological analysis

Flies were dissected to separate head and body without legs and wings. Immediately transferred into 4%PFA in phosphate buffered saline (PBS) for 15 minutes with mild agitation at room temperature. Later the tissues were washed with 1xPBS three times for 10 minutes each. Finally, tissues were transferred into 10% sucrose in PBS solution for overnight. The tissues were processed and prepared the slides according to standard protocol^29,41^. The tissues were stained with primary (Synapsin antibody 1:250 UI Developmental studies Hybridoma bank #3C11), phalloidin (Anova U0381) for overnight and washed. Secondary antibody AlexaFlour-750 anti-mouse 1:500 dilution added to the slides and incubated for 1 hour at room temperature. (Thermo Fisher #A-21037), for lipid analysis we used Lipid spot-488; 1:100 dilution (Biotinum #70065). Finally, we washed the slides three times with 1xPBS for 5 minutes each. We used VECTASHEILD Vibrance H-1800 which contains antifade mounting medium with DAPI (0.9ug/ml). The slides were left for overnight in dark and then images were captured using multichannel fluorescent microscopy (Olympus BX63) the images were analyzed using CellSens software at 10x magnification.

Region of interest (ROI) was defined using DAPI channel. As it distinguishes the anatomical subregions in head also in the body. Two ROIs were drawn for head and brain separately. One ROI drawn for body region. For each ROI, mean object area and object size were collected at 488 nm channel and mean pixel intensity was collected at 750 nm channel. From each group multiple sections were analyzed from head and body regions to measure the mean object count and mean object area. Lipid channel (488 nm) was thresholded to minimize background signal. The minimum detection size of object set to 20 pixels. Synapsin (750 nm) channel not thresholded to retain the integrity of comparison between groups and conditions. Essential background filters and thresholding values were applied before taking the images for all images across groups. To reduce the bias, we have collected data from multiple sections per fly (n=3) and multiple flies for each condition (n=5) for both head and body regions in all groups. Watershed segmentation and Otsu thresholding to the images were applied to all images for quantification. Sample processing, image capture and analysis performed using blinded genotype labelling to minimize bias.

### 2.7. Gene expression analysis

The heads and body tissues of the experimental flies were dissected and snap frozen in liquid N_2_ as previously reported^34,42^. The RNA was isolated using Zymo research Quick-RNA Microprep Kit (Catalog R1051, Zymo research Irvin California, USA). BioTek SYNERGY|LX multi-mode reader was used to assess RNA purity and quantity prior to cDNA synthesis for each biological replicate. cDNA was prepared using BioRad iScript™ Reverse Transcription Supermix (Catalog #170-8840). Quantitative PCR was conducted using the Sso advanced universal SYBR Green supermix from BioRad (Catalog #1725270) using BioRad CFX Opus Real-Time PCR system. No-template controls and melt-curve analysis used to confirm primer specificity and absence of contamination. Results represent 2^−ΔΔCt^ values normalized with RpL11 gene expression and control samples. All the qPCRs were performed using biological triplicates (n=3). Using GraphPad Prism 10 software Mean and standard deviation (SD) were calculated. The forward and reverse primers used for quantitative PCR are mentioned below. UPD1 FWD-CAGCGCACGTGAAATAGCAT; UPD1 REV-CGAGTCCTGAGGTAAGGGGA; UPD2 FWD-AGCGTCGTGATGCCATTCA; UPD2 REV-GCGATACGGATTGACATCGAA; UPD3 FWD-ATCCCCTGAAGCACCTACAGA; UPD3 REV-CAGTCCAGATGCGTACTGCTG; DOME FWD-CTCACGTCTCGACTGGGAAC; DOME REV-AGAATGGTGCTTGTCAGGCA; HOP FWD-CACCACCAACACCAATTC; HOP REV-GGAACGTCGTTTGGCCTTCT; STAT92E FWD-CCTCGGTATGGTCACACCC; STAT92E REV-TGCCAAACTCATTGAGGGACT; RPL11 FWD-CGATCTGGGCATCAAGTACGA; RPL11 REV-TTGCGCTTCCTGTGGTTCAC; EIGER FWD-GATGGTCTGGATTCCATTGC; EIGHR REV-TAGTCTGCGCCAACATCATC; IMD FWD-TCAGCGACCCAAACTACAATTC; IMD REV-TTGTCTGGACGTTACTGAGAGT; REAPER FWD-TGGCATTCTACATACCCGATCA; REAPER REV-CCAGGAATCTCCACTGTGACT; HID FWD-CACCGACCAAGTGCTATACG; HID REV-GGCGGATACTGGAAGATTTGC.

### 2.8. Microbiome analysis

Microbial quantification was performed using standardized protocol^31^ by washing the experimental flies with 70% ethanol and dissecting the limbs and head under sterile conditions to minimize the environmental contamination. The microbial DNA was isolated using DNeasy Ultraclean Microbial kit from Qiagen (catalog # 1222450). DNA was quantified using BioTek SYNERGY|LX multi-mode reader. Further we also run no template control test to further confirm possible environmental contamination. The DNA quantified and used 5 ng of pure DNA was used for each qPCR reaction using the primers listed below. Quantitative PCR was performed using the Sso advanced universal SYBR Green supermix from BioRad (Catalog #1725270) using BioRad CFX Opus Real-Time PCR system. Results represent 2^−ΔΔCt^ values normalized with RpL11 gene expression and control samples. 8-10 flies were used per replicate. All the qPCRs were performed using biological triplicates (n=3). Using GraphPad Prism 10 software Mean and SD were calculated. Primers used for qPCR analysis of microbiome are listed here, *Lactobacillus* (Genus) FWD-AGG TAA CGG CTC ACC ATG GC; REV-ATT CCC TAC TGC TGC CTC CC; *Lactobacillus brevis* FWD-GAC GTG CTT GCA CTG ATT TC; REV-CCG AAG CCA CCT TTC AAA C; *L. plantarum* FWD-CGA ACG AAC TCT GGT ATT GAT TG; REV-ACC ATG CGG TCC AAG TTG; *Acetobacter* (Genus) FWD-TAG TGG CGG ACG GGT GAG TA; REV-AAT CAA ACG CAG GCT CCT CC; *Acetobacter pasteur* FWD-CCGGCGGTGATCTTCTGTTC; REV-CCGCTCTGTGCGTCAAACTT; *A. promorum* FWD-CTA GAT GTT GGG TGA CTT AGT CA; REV-CGG GAA ACA AAC ATC TCT GCT TG; *Acetobacter pasteurianus* FWD-CCGGCGGTGATCTTCTGTTC; REV-CCGCTCTGTGCGTCAAACTT; 16s rRNA universal FWD-AGAGTTTGATCCTGGCTCAG; REV-GGTTACCTTGTTACGACTT; RPL11 FWD-CGATCTGGGCATCAAGTACGA; REV-TTGCGCTTCCTGTGGTTCAC.

### 2.9. Western blot analysis

Western blot analysis was performed^43^ using a standard protocol. The experimental flies with appropriate controls were dissected to separate limbs then head and body tissues. The head tissues were added with RIPA buffer (Thermo scientific, Pierce^TM^ RIPA buffer) provided with protease and phosphatase inhibitor (Thermo scientific, Halt^TM^ catalog **#**78441) in 1.5 ml Eppendorf tube. The heads were gently crushed and then homogenized with a handheld homogenizer (Fisher brand motorized tissue grinder 12-1413-61) in ice. The homogenized tissue samples were centrifuged at 16000 rpm for 15 minutes at 4 °C. The supernatant was removed carefully without disturbing pellet. Protein quantity was estimated using Bradford reagent (BioRad catalog #5000006) at 595 nm the emission was measured using microplate reader (BioTek, SYNERGY multimode reader). Samples were prepared by using 2x sample buffer (BioRad catalog #1610737) and boiled at 100°C for 5 minutes. Samples were allowed to cool at room temperature and centrifuged briefly. Using precast mini gels (BioRad catalog #4561036) we ran the SDS-PAGE to separate the proteins according to their molecular weight. BioRad tri color protein marker (BioRad catalog #1610374) was used as molecular weight marker. The gels were subjected to transfer overnight on to Nitro cellulose membrane 0.2 um (BioRad catalog # 1620112) overnight at 4 °C at 25 volts using transfer buffer with 20% methanol. The blots were briefly washed with 1x TBS and blocked with 3% BSA (Fisher scientific) for one hour at room temperature and washed the excess blocking buffer using TBS briefly. Primary antibodies Tau (DSHB, 5A6-C), phospho tau Ser 202, Thr205 (Invitrogen, MN1020), amyloid beta 42 (Sigma Aldrich, AB1510), Synaptotagmin (DSHB 3H2 2D7), Ref2p (Ab178440), ubiquitin (Abcam-EP8589), AMPK (Ab80039), pAMPK (Abcam-EPR5683), GABARAP (CST-E1J4E#13733), α-tubulin (DSHB 4A1) in TBST and the blots were incubated overnight at 4 °C. Next day blots were washed with TBS-TBST-TBS for 30 minutes 10 minutes each, then added secondary anti-rabbit (Abcam #ab6721), or anti mouse (Abcam #ab6728) tagged with HRP and incubated at room temperature for one hour. After secondary antibody treatment blots were washed with TBST then TBS followed by TBST 10 minutes each for 30 minutes to remove excess secondary antibody. ECL reagents (BioRad) were used to develop the blots using Amersham600 chemidoc with chemiluminometric reading. To confirm the equal protein loading across the sample’s normalization was performed using α-tubulin. Chemiluminescence exposures used within linear detection range. Quantification of blots were performed using Image J and used similar quantification parameters across all experimental groups. 8-10 heads and body tissues were used in each replicate (n=3).

## 3. Statistical analysis

Statistical analyses were carried out using GraphPad Prism version 10. Data from each overexpression gene was compared with the control (*Glaz*/+) and *Glaz/GFP* (provided in supplementary file) in genotypes and with or without microbiome. Data are reported as mean ± SD. Statistical significance was defined as the following: p < 0.05 (*), p < 0.01 (**), p < 0.001(***), p < 0.0001 (****). Outlier test is performed using modified Z-score method^44^. Detailed statistical analyses among different genotypes in conventional control and axenic flies have been shown in the source data. Moreover, as indicated under each figure legends section, we used one-way ANOVA followed by Dunnett’s post hoc test, for microbiome analysis. Two-way ANOVA followed by Sidak’s multiple comparisons test for immunofluorescence, qPCR, western blot analysis memory assay and geotaxis analysis. chi-square test is also used for memory assay (reference). Sample size, randomization and details about biological replicates were mentioned in the figure legends separately.

## 4. Results

### 4.1. Alzheimer’s disease pathology alters Gut-microbiome composition

To understand whether gut microbiome is distinct in AD and how different AD related genetic mutations affect the microbiome and pattern of disease progression, we have used humanized transgenic *Drosophila* model carrying AD related genetic mutations in tau protein (referred as HTau, HTau*^R406W^*, phospho-HTau), amyloid precursor protein (APP), and a*β*42 coding transgenes. All these trans genes were expressed under glial cell specific driver (*Glaz-GAL4*). We have checked the changes in the quantity of most abundant bacterial population in the gut, i.e., *Lactobacillus* and *Acetobacter* in our AD fly models using real time PCR. *Lactobacillus* genus (Figure. 1D), found to be increased in, phospho HTau (p<0.0001, 5.19±1.78), a*β*42 (p<0.0001, 6.89±0.87) and APP (p<0.0001, 11.85±1.76) compared to control (*Glaz/+*) flies (hereafter the control represents *Glaz*/+). Species specific analysis revealed that *L. brevis* (Figure. 1E) abundance in all the AD fly models HTau (p<0.03, 0.43±0.16), HTau*^R406W^* (p<0.03, 0.44±0.06), phospho HTau (p<0.0019, 0.55±0.058), and APP (p<0.0001, 0.705±0.33) and *L. plantarum* (Figure. 1F) increased HTau (p<0.0045, 3.95±1.30), HTau*^R406W^* (p<0.0061, 3.86±1.02), phospho HTau (p<0.0001, 8.21±1.45), a*β*42 (p<0.0001, 5.75±1.14) and APP (p<0.0001, 9.58±2.76) significantly. Whereas *Acetobacter* (Figure. 1G) abundance showed significant increase in HTau (p<0.0045, 3.31±0.37), phospho HTau (p<0.0001, 4.10±0.311), a*β*42 (p<0.0027, 1.83±0.652), and in APP (p<0.0001, 2.29±0.60) compared to control. *A. promorum* (Figure. 1H) showed significant increase in HTau (p<0.0002, 2.82±1.14) with no significant change in other AD models. *A. pasteurianus* (Figure. 1I) showed similar trend in increase like *Acetobacter genus* in HTau (p<0.0001, 7.51±1.35), phospho HTau (p<0.0001, 5.43±1.25), a*β*42 (p<0.0001, 3.58±0.67), and in APP (p<0.0001, 3.69±0.48) compared to control. Since the microbiome abundance varies significantly among the AD transgenic fly models, we further aimed to understand the impact of complete loss of gut microbiome on AD pathology. To prepare microbiome free AD flies, we administered antibiotic cocktail (ampicillin, tetracycline, kanamycin, and erythromycin) supplemented food after dechorionizing the embryos for up to 3-4 weeks for our study. We then quantified the microbiome abundance using universal 16s rRNA primers, in conventional control, (here after termed as CC) to understand the microbiome abundance (Figure. 1C) in HTau (p<0.0001, 6.78±1.76), phospho HTau (p<0.0001, 9.95±1.77), a*β*42 (p<0.0001, 2.45±0.53), and in APP (p<0.0001, 10.37±2.59) compared to control. In axenic antibiotic flies (here after termed as AA) (Figure. 1B), we found ∼90% reduction in microbial population in AA flies *Glaz/+* (p<0.0001, 0.07±0.01), GFP (p<0.0001, 0.01±0.003), HTau (p<0.0001, 0.047±0.039), HTau*^R406W^*(p<0.0001, 0.007±0.001), phospho HTau (p<0.0001, 0.029±0.0146), a*β*42 (p<0.0001, 0.015±0.006), and in APP (p<0.0001, 0.046±0.018) compared to CC control *Glaz/+* (0.812±0.24) flies (SI. Figure. 1A-H). This provides us with a model system to test the impact of microbiome on behavioral and metabolic phenotypes during AD pathology. We used this model to understand the impact of gut microbiome on sleep/circadian behavior, cognitive/memory functions, locomotory activity. Further we measured how it influences lipid accumulation, protein aggregation, neuroinflammation and cell death during AD pathology both under tauopathy and amyloid deposition.

**Figure 1.**
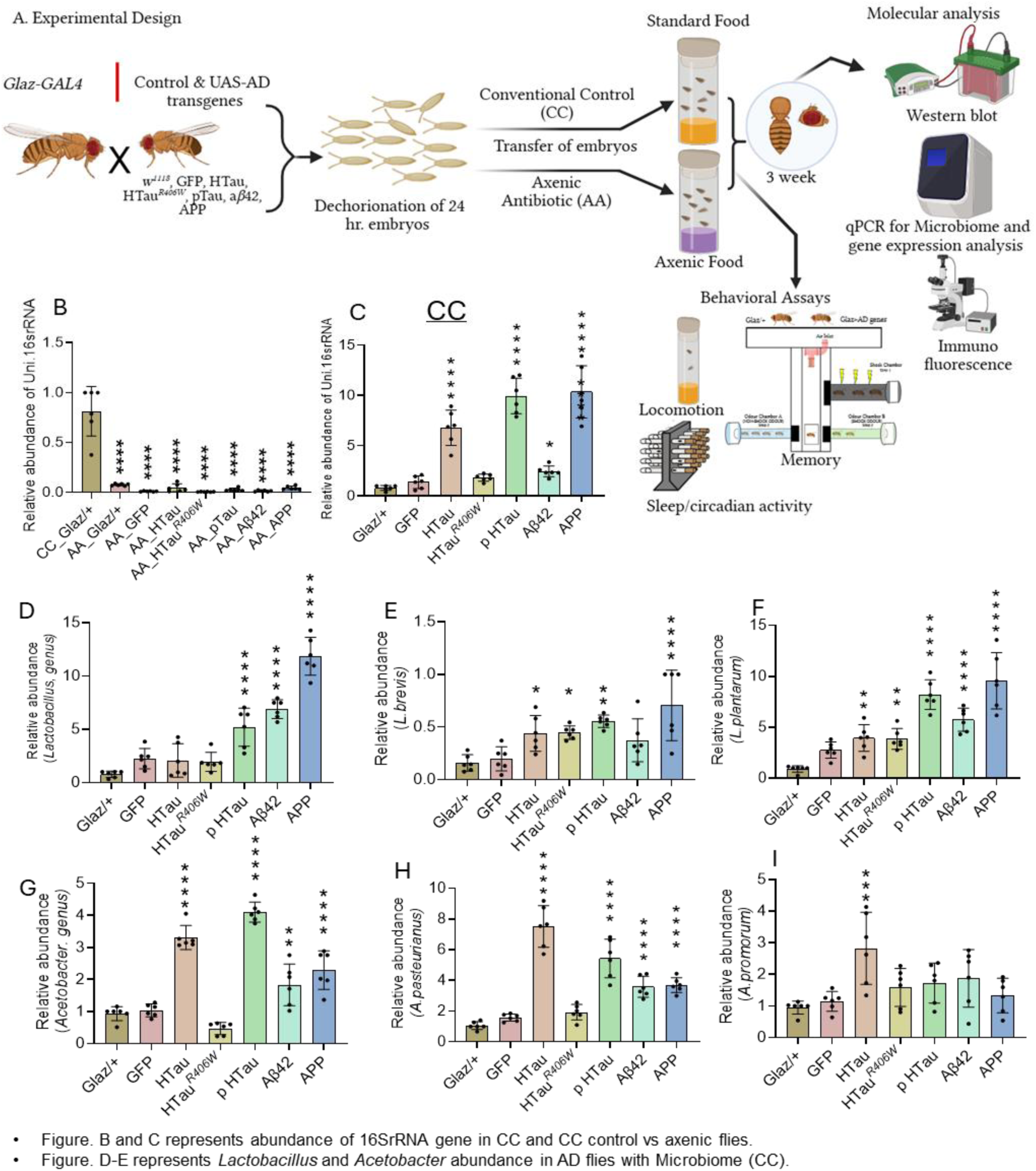
1A. Experimental design, representing the driver line and experimental fly lines, embryo collection time points, washing and antibiotics (Erythromycin, Kanamycin, Tetracycline, Ampicillin) added to the standard fly food (axenic antibiotic-AA) and standard food (conventional control-CC) aging and microbiome quantification procedures. Gut microbiome showed significant alterations among humanized AD models. Each figure represents quantification through qPCR, (B). Comparison of microbiome abundance between *Glaz*/+ of conventional control (CC) group and all axenic antibiotic (AA) group flies shows 90% reduction (p<0.0001) in AA flies gut microbiome abundance. (C). Represents the abundance of gut microbiome in all flies measured using universal 16srRNA primers in CC flies, we found a significant increase in HTau, phospho HTau, APP (p<0.0001) and a*β*42 (p<0.05). (D). *Lactobacillus* genus abundance in control and AD flies’ body, expressing genes under glial cell specific driver (*Glaz-GAL4*) at 3 weeks age. Among them phospho HTau and APP (p<0.0001) showed a significant increase. (E). *L. brevis*, increased in HTau, HTau*^R406W^*, a*β*42 (p<0.05), phospho HTau (p<0.01) and APP (p<0.0001), (F). *L. plantarum*, showed increased abundance in HTau*^R406W^*(p<0.01), phospho HTau, a*β*42, APP (p<0.0001) flies. (G). *Acetobacter* genus, showed decrease in HTau*^R406W^* (p<0.01), increased in phospho HTau, APP (p<0.0001), and a*β*42 (p<0.01) flies. (H). *A. promorum* showed increase in HTau (p<0.001), and no significant change in other genotypes. (I). *A. pasteurianus* no significant difference in HTau*^R406W^,* and increase in HTau, phospho HTau, a*β*42, and APP (p<0.0001) All the microbiome quantifications were compared with control (*Glaz*/+) group in B-I. For microbiome quantification experiment One-way ANOVA was used because this panel compares multiple genotypes within a single condition. Dunnett’s post hoc test was used as comparisons drawn with single control group. n=3, and p-values * <0.05, ** <0.01, *** < 0.001, **** <0.0001.

### 4.2. Gut microbiome influencing lipid metabolism and synaptic levels in the brain

The lipid object counts in head (Figure. 2B) showed no difference among AD genotypes in CC flies but found significant increase in HTau*^R406W^* (p<0.0001, 257.13±76.11) and phospho HTau (p<0.012, 221.93±73.45) compared to AA control (*Glaz/+*) and CC HTau*^R406W^* (p<0.0001, 141.53±45.19) and phospho HTau (p<0.0001, 115.48±26.23) flies. Lipid object area in head (Figure. 2C) showed significant reduction in AA GFP (p<0.0001, 272±166.95), HTau (p<0.0075, 441.40±101.83), HTau*^R406W^*(p<0.0001, 298.95±153.70), phospho HTau (p<0.0001, 267.22±73.53), a*β*42 (p<0.0001, 300.32±107.31), and in APP (p<0.0001, 301.43±147.21) compared to control (651.91±231.43) and decreased even with CC GFP (665.65±192.89) and a*β*42 (512.23±145.72) flies. In the brain region object count (Figure. 2D) was increased in APP flies (p<0.0001, 56.37±24.23) compared to control (12.50±7.17) in CC, and HTau (p<0.0001, 54.31±38.54), phospho HTau (p<0.01, 23.35±9.27) in AA flies. We also observed significant reduction in APP flies (p<0.0001, 6.25±4.18) in AA condition compared to CC, APP flies. Lipid object area in the brain (Figure. 2E) was reduced in all genotypes in AA compared to controls HTau*^R406W^* (p<0.0001, 98.57±68.45), phospho HTau (p<0.001, 151.99±107.53), a*β*42 (p<0.0001, 52.24±40.59), and in APP (p<0.0001, 155.85±132.76) and no significant change compared to CC flies. In the abdomen region lipid object count (Figure. 2I) did not show significant difference in CC flies compared to controls and in AA flies only HTau*^R406W^* (p<0.016, 479.68±137.78) and APP flies (p<0.046, 503.76±150.75) showed significant increase compared to AA control (274.57±46.79). The lipid object area (Figure. 2J) showed significant increase in HTau (p<0.003, 512.65±154.75), phospho HTau (p<0.0001, 579.6±138.99), and a*β*42 (p<0.034, 472.83±141.98) in CC flies. Whereas in AA flies phospho HTau (p<0.0001, 608.94±105.43) showed significant increase and APP (p<0.001, 170.62±100.09) showed significant decrease compared to AA control and GFP (p<0.0001, 215.2±161.66) showed significant reduction compared to AA control (370.72±129.39) and CC (354.95±90.96) flies.

**Figure 2.**
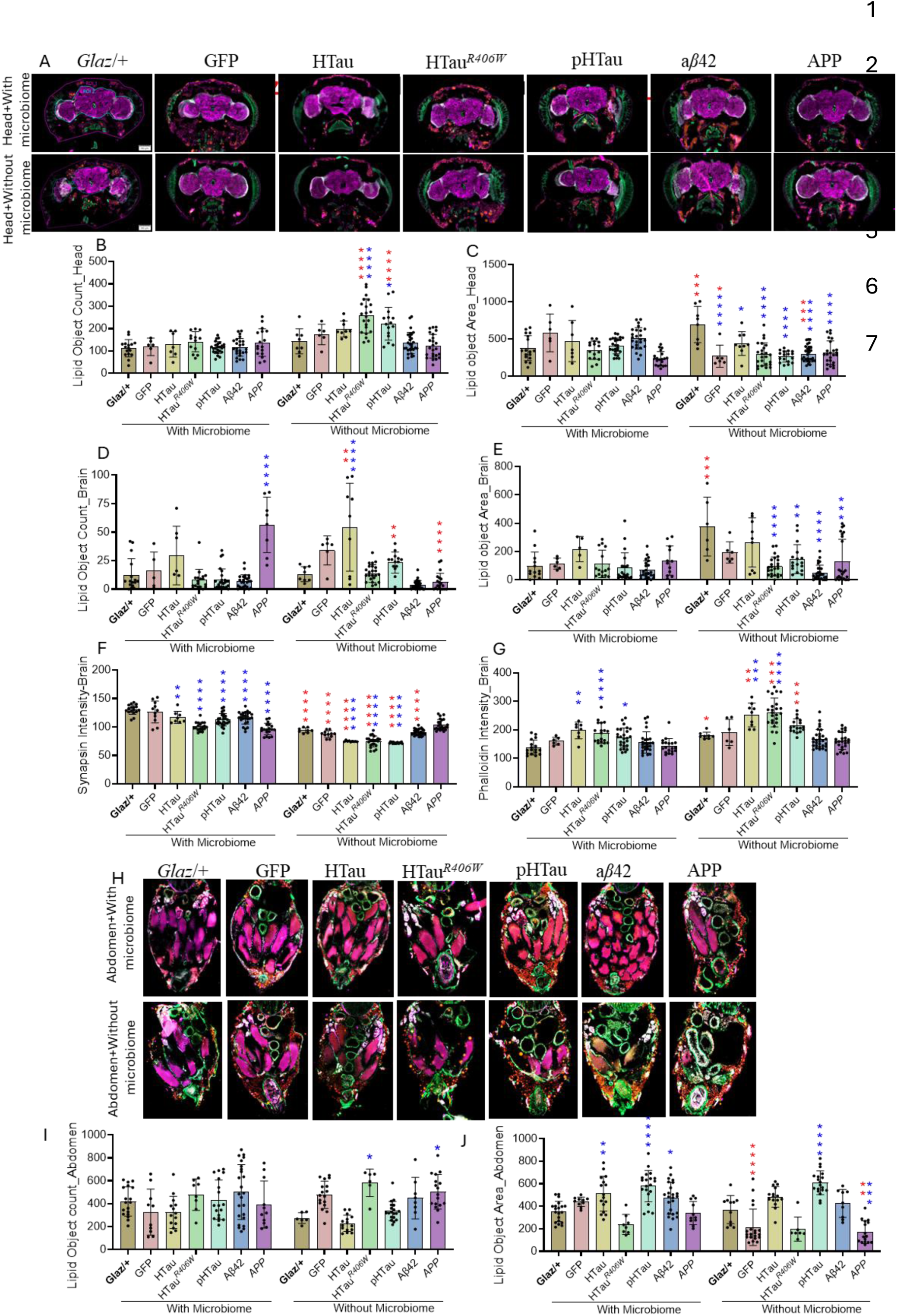
Depletion of microbiome regulates lipid metabolism and synaptic integrity. Immunofluorescence analysis for CC, AA, AD transgenes expressing flies and control (A-G) head and (H-J) abdomen. (A). Staining in heads, (B). staining in abdomen regions, (C). Lipid object counts in head, in CC no difference observed, and in AA flies increase in HTau*^R406W^* (p<0.0001), and phospho HTau (p<0.05) observed compared to controls. Also compared to CC flies AA HTau*^R406W^*, and phospho HTau (p<0.0001) showed increased lipid count in heads. (D). Lipid object area in head, AA flies showed significant decrease in GFP, HTau*^R406W^*, phospho HTau, a*β*42 and APP (p<0.0001), and in HTau (p<0.05) compared to control. Whereas compared to CC flies, AA flies showed reduced lipid object area in GFP (p<0.05), a*β*42 (p<0.001), and increased in *Glaz*/+ (p<0.001) observed. (E). Lipid object counts in brain, increased in HTau (p<0.05) and APP (p<0.0001) in CC flies, and GFP (p<0.05), HTau (p<0.0001) in AA flies compared to controls. Compared to CC flies HTau and phospho HTau (p<0.01) showed increased and decreased in APP (p<0.0001) lipid object count in brain was observed. (F). Lipid object area in brain, CC flies no change, but AA flies HTau*^R406W^*, a*β*42 (p<0.0001), APP (p<0.001) and phospho HTau (p<0.01) significant rection compared to control. In *Glaz*/+ (p<0.001) significant increase observed in AA flies compared to CC flies. (G). Synapsin intensity in brain, reduced in all AD flies significantly GFP (p<0.05), HTau (p<0.01) and HTau*^R406W^*, phospho HTau, a*β*42, and APP (p<0.0001) and in AA flies HTau, HTau*^R406W^*, and phospho HTau showed a significant reduction (p<0.0001) compared to controls. But in AA condition all the flies showed reduced synapsin compared to CC flies *Glaz*/+, GFP, HTau*^R406W^*, phospho HTau, a*β*42, and APP (p<0.0001) compared to CC flies. (H). Phalloidin intensity in brain, increased in HTau (p<0.001), HTau*^R406W^*(p<0.0001), phospho HTau (p<0.01), and a*β*42 (p<0.05) in CC flies, and HTau, HTau*^R406W^* (p<0.0001) showed significant increase compared to controls. Compared to CC flies, AA flies showed increase in *Glaz*/+ (p<0.05), HTau (p<0.01), and HTau*^R406W^*, phospho HTau (p<0.001) phalloidin intensity observed. (I) Lipid object counts in abdomen, observed significant increase only in HTau*^R406W^*, APP (p<0.05) flies in AA condition compared to control. (J). Lipid object area in abdomen increased in HTau (p<0.01), phospho HTau 9p<0.0001), and a*β*42 (p<0.05) in CC flies and in AA flies increased in HTau (p<0.05), HTau*^R406W^* (p<0.01) compared to control, and reduced in APP (p<0.05) compared to both control and APP flies in CC. Two-way ANOVA to establish significance, followed by Sidak’s multiple comparisons test to assess the effects of genotype and microbiome condition. n=3, and p-values * <0.05, ** <0.01, *** < 0.001, **** <0.0001.

Since AD patients showed significant increase in lipid deposition and lower synaptic level that contributed to memory disfunction^45,46^. We have checked the synapsin expression (Figure. 2F) in heads of both CC and AA flies expressing AD genes. We found a significant reduction of synapsin HTau (p<0.01, 113.81±6.35), HTau*^R406W^*(p<0.0001, 100.8±7.66), phospho HTau (p<0.0001, 112.44±11.06), a*β*42 (p<0.0001, 116.17±10.71), and in APP (p<0.0001, 96.33±10.42) flies compared to control (130.40±8.62)and in AA flies we have observed a further reduction of synapsin in control (p<0.0001, 93.82±4.51), GFP (p<0.0001, 87.24±6.3), HTau (p<0.0001, 74.53±0.61), HTau*^R406W^*(p<0.0001, 73.79±9.33), phospho HTau (p<0.0001, 72.16±1.24), and a*β*42 (p<0.0001, 90.77±5.13) compared to CC flies. Whereas in AA flies, HTau, HTau*^R406W^*, phospho HTau showed (p<0.0001) further reduction of synapsin compared to AA control flies. We then also checked phalloidin staining (Figure. 2G) in AD flies, we have observed a significant increase in HTau (p<0.0014, 199.72±32.81), HTau*^R406W^* (p<0.0003, 182.89±30.57) and phospho HTau (p<0.031, 172.13±35.38) in CC compared to control (139.94±27.21) and HTau (p<0.0005, 251.99±45.79) and HTau*^R406W^* (p<0.0001, 267.29±48.42) in AA compared to control flies (177.61±5.89). In AA flies we further observed an increase in phalloidin staining HTau (p<0.002), HTau*^R406W^* (p<0.0003) and phospho HTau (p<0.027) compared to CC flies. In both CC as well as AA flies we did not observe any significant difference in phalloidin staining between CC and AA control flies. Our results suggest that AD-related genetic mutations in AA flies altering lipid droplet accumulation dynamics across head, brain and abdomen regions, compared to CC flies. Genotypic specific impact of phalloidin staining suggests possible actin remodeling and reduced synapsin levels further support the influence of microbiome on synaptic health during AD.

### 4.3. Depletion of microbiome leads to enhanced inflammatory responses in the head and body regions of humanized AD model

To evaluate the impact of axenic conditions on inflammatory responses in AD flies we have tested the inflammatory gene expression pattern during AA and CC condition (Figure. 3) in both heads (Figure. 3A-E) and body (Figure. 3F-J) regions using qPCR. We have tested the cytokine *Upd3* (IL6) in heads of CC flies where we found decrease in GFP (p<0.03, 0.16±0.10) and increase in APP (p<0.03, 1.88±0.18) compared to control (0.98±0.14) flies. In AA flies, HTau*^R406W^* (p<0.0003, 2.29±0.54), a*β*42 (p<0.0001, 2.7±0.41) and APP (p<0.0001, 2.58±0.41) flies showed significant increase in *Upd3* expression compared to control and significant increase in HTau*^R406W^*(p<0.0001), a*β*42 (p<0.0001) in AA compared to CC flies. The UPD3 cytokine receptor gene *Dome* (Figure. 3B) showed significant increase in a*β*42 (p<0.0035, 2.66±0.21) and APP (p<0.04, 1.94±0.42) in CC files compared to controls (0.988±0.07). In AA flies HTau*^R406W^* (p<0.0018, 2.73±0.18), a*β*42 (p<0.0001, 4.41±0.58) and APP flies (p<0.0001, 4.25±1.28) showed significant increase compared to AA control (0.966±0.021) flies. Compared to CC flies AA, HTau*^R406W^*(p<0.0032), a*β*42 (p<0.0001) and APP flies (p<0.0001) showed significant increase and phospho HTau (p<0.048, 1.95±0.41) showed significant decrease. The cytoplasmic downstream target of DOME known as *Hop* showed significant increase in a*β*42 (p<0.0024, 2.27±0.24) and APP (p<0.0001, 2.89±0.58) in CC files compared to controls (0.93±0.07). In AA flies HTau*^R406W^* (p<0.0001, 4.55±0.62), a*β*42 (p<0.0001, 3.59±0.24) and APP flies (p<0.0001, 4.63±0.45) showed significant increase compared to AA control (1.07±0.11) flies. Further, compared to CC flies AA, HTau*^R406W^*(p<0.00012), a*β*42 (p<0.0028) and APP flies (p<0.0001) showed significant increase. Up on ligand binding (*Upd*3), HOP phosphorylates DOME and then activates STAT92E, the downstream target of HOP^47,48^. In CC flies *Stat92e* expression (Figure. 3D) has no significant difference except GFP (p<0.036, 0.91 ±0.079) showed decrease expression compared to control (0.94±0.084). In AA condition HTau*^R406W^* (p<0.004, 1.45 ±0.09) phospho HTau (p<0.0003, 1.69±0.15), and APP flies (p<0.0004, 1.82 ±0.18) showed significant upregulation of *Stat92e* compared to AA control (0.91±0.07) flies. Further, compared to CC flies AA, HTau*^R406W^*(p<0.001), a*β*42 (p<0.003) and APP flies (p<0.014) showed significant increase in *Stat92e* gene expression. We further checked the possible implication of inflammation on promoting the cell death. We have checked the *Hid* which is an inhibitor of *Drosophila* apoptosis inhibitory protein1 (DIAP1)^49^. In our study we found a significant increase in the HTau*^R406W^* (p<0.03, 1.60±0.34), and phospho HTau (p<0.0001, 2.41±0.24) in AA flies compared to AA control (0.99±0.06) flies. Further, compared to CC flies AA, HTau*^R406W^*(p<0.0001), phospho HTau (p<0.0001) and APP flies (p<0.0009) showed significant increase in *Hid* expression. In CC flies we did not observe any significant change in *Hid* expression compared to control. We further checked *Upd*3 (Figure. 3F), *Dome* (Figure. 3G)*, Hop* (Figure. 3H)*, Stat92e* (Figure. 3I) and *Hid* (Figure. 3J) in abdomen region to understand how absence of gut microbiome influences peripheral inflammatory responses. In our study we haven’t observed any difference among CC flies in *Upd*3 expression whereas in AA, HTau (p<0.0001, 2.66±0.32), phospho HTau (p<0.008, 0.95±0.30), a*β*42 (p<0.0001, 2.43±0.061) and APP (p<0.0001, 2.46±0.55) showed significant increase compared to AA control (0.94±0.12). Further, compared to CC flies AA, HTau (p<0.0001), phospho HTau (p<0.007), a*β*42 (p<0.0001) and APP flies (p<0.0001) showed significant increase in *Upd3* expression. *Dome* showed decreased expression in CC, HTau*^R406W^* (p<0.0016, 0. A significant increase in AA, phospho HTau (p<0.0001, 2.06±0.23), a*β*42 (p<0.0001, 1.911±0.17), and APP (p<0.0001, 2.44±0.29) observed compared to AA, control (0.99±0.011). Further, increase in *Dome* expression observed in AA, HTau*^R406W^*(p<0.0009, 0.99±0.01), phospho HTau (p<0.0002), a*β*42 (p<0.0001) and APP flies (p<0.0001) showed compared to CC flies. In CC flies *Hop* expression increased only in phospho HTau (p<0.04, 1.52±0.11), whereas in AA flies phospho HTau (p<0.0001, 2.19±0.03), a*β*42 (p<0.0001, 1.87±0.03), and APP (p<0.0001, 0.6±0.13) showed significant increase compared to AA (0.97±0.06) and CC (0.95±0.007) controls. *Stat92e* showed no change in CC flies, but in AA flies HTau*^R406W^* (p<0.02, 1.49±0.21) and APP (p<0.0001, 2.31±0.16) flies showed significant increase compared to control (1.012±0.02). Compared to CC flies in AA flies HTau (p<0.001), HTau*^R406W^*(p<0.0001), and APP (p<0.0001) showed a significant increase in *Stat92e* expression. Further, we have observed a significant increase in cell death marker expression in AA condition in phospho HTau (p<0.0001, 1.99±0.19), a*β*42 (p<0.003, 1.59±0.25), and APP (p<0.0003, 1.73±0.10) compared to AA control (0.94±0.06). Compared to CC flies HTau, (p<0.0008), phospho HTau (p<0.0002), a*β*42 (p<0.0001) and APP (p<0.0001) showed a significant increase in AA flies. We also checked the *Upd1* (SI. Figure. 3A)*, Upd2* (SI. Figure. 3B) cytokine genes, *Eiger* (SI. Figure. 3C)*, Imd* (SI. Figure. 3D) involved in antimicrobial peptide (AMPs) production. We further checked another cell death marker *Reaper* (SI. Figure. 3E). In all these genes HTau*^R406W^*and APP showed significant upregulation in head regions. In the body *Upd1* expression decreased (SI. Figure. 3G) in HTau, phospho HTau, a*β*42 and APP in CC flies whereas in AA condition, these flies showed significant upregulation compared to control and CC flies. *Upd2* (SI. Figure. 3H) HTau showed reduction in CC flies compared to control and in AA flies HTau, a*β*42 and APP showed significant upregulation compared to control and CC flies. In CC flies *Eiger* (SI. Figure. 3I), *Imd* (SI. Figure. 3J) and *Reaper* (SI. Figure. 3K) gene expression did not show any significant change, whereas in AA flies except HTau*^R406W^* for *Eiger* and a*β*42 for *Imd* all the AD genes showed a significant upregulation in AA condition compared to control and CC flies. The SI. Figure. 3F shows the molecular communication between cytokines and *Stat92e* for inflammatory gene expression. Based on our study we can conclude that the axenic condition significantly amplifies the inflammatory response in AD flies with the upregulation of *Upd*3, *Dome*, *Hop*, and *Stat92e* compared to conventional control flies. It is further accompanied by upregulated pro-apoptotic markers such as *Hid* and *Reaper* involved in cell death marker gene expression. Even though microbial metabolites were not directly measured in our study, the increased *Upd3–Dome–Hop–Stat92e* signaling in AA flies suggesting a possible microbiome-sensitive inflammatory axis in glial AD models.

**Figure 3.**
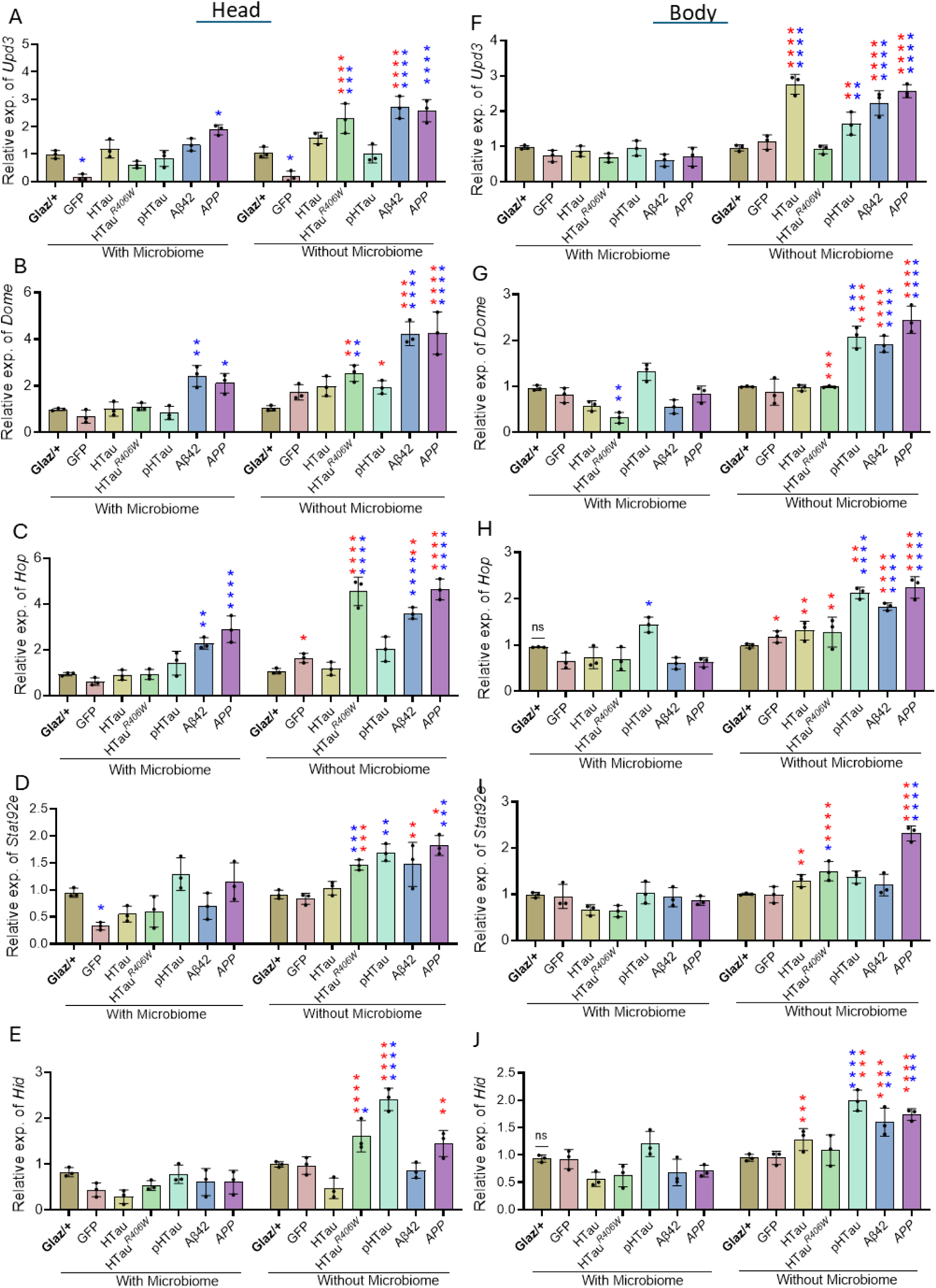
Depletion of microbiome further enhanced AD linked inflammation in the body and brain regions in humanized AD model. Changes in inflammatory gene expression patterns in CC and AA flies in the head (A-D) and in body (F-I) at 3 weeks age quantified using qPCR. *Upd*3 reduced in GFP(p<0.05) ad increased in APP (0.05) in CC flies and reduced in AA flies in GFP (p<0.05) and increased in HTau*^R406W^*(p<0.001), a*β*42 and APP (p<0.0001) compared to controls. In AA flies HTau*^R406W^*, a*β*42 (p<0.001) showed significant increase compared to CC flies, (B). *Dome* increased in CC flies in a*β*42 (p<0.01) and APP (p<0.05), in AA flies HTau*^R406W^* (p<0.01) and in a*β*42, APP (p<0.0001) compared to control. In addition, compared to CC flies AA showed significant increase in HTau*^R406W^* (p<0.01), phospho HTau (p<0.05), a*β*42 (p<0.01), and APP (p<0.0001) compared to CC flies. (C). *Hop* increased in a*β*42 (p<0.01) and APP (p<0.0001), in AA flies HTau*^R406W^*, a*β*42, and APP (p<0.0001) compared to control. Compared to CC flies GFP (p<0.05), HTau*^R406W^*, and APP (p<0.0001) and a*β*42 (p<0.01) in AA flies increased in *Hop.* (D). *Stat92e* showed significant reduction in GFP (p<0.05) in CC flies and in AA flies significant increase in HTau*^R406W^*, APP (p<0.001) and phospho HTau (p<0.01) compared to control. In AA flies a significant increase in HTau*^R406W^* (p<0.001), a*β*42 (p<0.01), and APP (p<0.05) compared to CC flies. (Figure. E). Cell death marker *Hid* significant reduction in HTau (p<0.05), CC flies, and significant increase in HTau*^R406W^* (p<0.05) and phospho HTau (p<0.0001) in AA flies compared to control. Also, AA flies showed increased *Hid* in HTau*^R406W^*, phospho HTau (p<0.0001) and APP (p<0.01) compared to CC flies in the head. (F). In the body Upd3 has no difference in CC flies, but in AA flies HTau, a*β*42, APP (p<0.0001), and in phospho HTau (p<0.01) showed significant increase compared to controls. (G). *Dome* reduced in HTau*^R406W^* (p<0.01) in CC flies and increased in phospho HTau (p<0.001), a*β*42, and APP (p<0.0001) compared to controls. Compared to CC flies HTau*^R406W^* (p<0.001), phospho HTau, a*β*42, and APP (p<0.0001) in AA flies’ shoed significant increase. (H). *Hop* increased in phospho HTau (p<0.05) in CC flies, and in AA flies phospho HTau (p<0.01), a*β*42, and APP (p<0.0001) showed significant increase in *Hop* compared to controls. Also, GFP (p<0.05), HTau, HTau*^R406W^*, phospho HTau (p<0.01), and a*β*42, and APP (p<0.0001) in AA flies compared to CC flies. (I). *Stat92e* no change observed in CC flies, whereas in AA flies significant increase in HTau*^R406W^*, phospho HTau (p<0.05), and APP (p<0.0001) observed. In addition, Hatu (p<0.01), HTau*^R406W^*and APP (p<0.0001) showed significant increase compared to CC flies in *Stat92e*. (Figure. J). *Hid,* has no significant change in CC flies but in AA phospho HTau (p<0.0001), a*β*42 (p<0.01), and APP (p<0.001) increased compared to controls. Also, AA HTau, phospho HTau (P<0.001) and a*β*42, APP (p<0.0001) flies showed significant increase in *Hid* compared to CC flies in the body. Two-way ANOVA to establish significance, followed by Sidak’s multiple comparisons test to assess the effects of genotype and microbiome condition. n=3, and p-values * <0.05, ** <0.01, *** < 0.001, **** <0.0001.

### 4.4. Depletion of microbiome further impaired sleep-wake cycle/circadian homeostasis linked with humanized AD model

In our study AD transgenic flies showed increased day activity (Figure. 4B) in GFP (p<0.0001, 990.68±133.10) phospho HTau (p<0.0001, 794.96±160.19) a*β*42 (p<0.0001, 273.46±94.34) and APP (p<0.0001, 829.65±302.06) during CC condition compared to control (539.74±251.64). In AA flies *Glaz/+* (p<0.0001, 840.39±128.01) HTau*^R406W^* (p<0.0001, 913.96±192.17) and a*β*42 (p<0.0071, 457.85±138.13) flies showed increased and GFP (p<0.0001, 570.60±81.16), phospho HTau (p<0.0001, 692.81±161.80) and APP (p<0.0001, 463.73±99.27) flies showed reduce day activity compared to CC flies. Whereas except HTau, and HTau*^R406W^* all the AA flies showed reduced (p<0.0001) day activity compared to AA control (*Glaz*/+) flies. Night activity increased in GFP (p<0.0001, 775.19±126.59), HTau (p<0.007, 575.70±246.53), HTau*^R406W^* (p<0.031, 561.13±206.15), and APP (p<0.0008, 554.56±246.09) flies (Figure. 4C) compared to CC control (437.27±190.15). In AA night activity decreased in HTau (p<0.0014, 467.73±122.12), phospho HTau (p<0.006, 491.37±182.44), a*β*42 (p<0.0001, 405.16±123.34), and APP (p<0.0085, 481.41±173.79), compared to control (675.71±136.61) flies. Total activity (Figure. 4D) increased in GFP (p<0.0001,1815.84±248.37), APP (p<0.0001, 1461.31±362.26) and reduced in a*β*42 (p<0.015, 774.46±134.37) flies compared to control (1072.50±361.95) in CC flies. In AA flies HTau (p<0.0009, 1191.32±309.79), phospho HTau (p<0.0001, 957.28±220.39), a*β*42 (p<0.0001, 881.41±1298.57) and APP (p<0.0001, 1074.51±349.59) flies showed reduced total activity compared to control (481.41±173.79). In AA flies’ control (p<0.0001) and HTau*^R406W^* (p<0.0001 1573.91±469.32) flies showed increased and GFP (p<0.0001, 1304.63±285.33), APP (p<0.0001) flies showed reduced total activity (Figure. 4D) compared to CC flies. Day sleep (Figure. 4E) reduced in GFP (p<0.0001, 354.16±36.58), phospho HTau (p<0.0001, 353.96±71.35) and APP (p<0.0001, 330.58±59.01), increased in a*β*42 (p<0.0001, 573.72±61.96) flies compared to control (464.93±89.89) in CC flies. In AA flies’ control (p<0.0008, 374.14±50.62), HTau*^R406W^* (p<0.0001, 155.44±73.79) and a*β*42 (p<0.0001, 407.74±73.18) showed reduced, and APP (p<0.0001, 498.45±56.64) showed increased day sleep compared to CC flies. In AA flies APP (p<0.0001) showed increased and HTau*^R406W^*(p<0.0001) showed reduced day sleep compared to AA control flies with no significant difference in HTau and phospho HTau flies. Night sleep (Figure. 4F) reduced significantly in GFP (p<0.028, 445.87±54.66), HTau (p<0.015, 453.75±101.71) HTau*^R406W^* (p<0.025, 457.64±93.58) and APP (p<0.046, 461.31±97.90) flies except in phospho HTau and a*β*42 flies compared to control (532.63±86.68) flies. In AA flies’ control (p<0.03, 459.12±61.02), GFP (p<0.0001, 333.93±86.8), HTau*^R406W^* (p<0.0001, 316.42±120.98) and phospho HTau (p<0.016, 486.16±101.64) showed reduced night sleep compared to CC flies. Among the AA fly’s night sleep reduced in GFP (p<0.0008), HTau*^R406W^*(p<0.0001, 316.42±120.98) and increased in a*β*42 (p<0.021, 537.17±65.31) flies compared to control. Total sleep (Figure. 4G) reduced in GFP (p<0.0004, 782.34±78.19) and APP (p<0.0001, 797.12±134.69) compared to control in CC (976.72±194.30) flies. In AA flies’ control (p<0.0041, 833.14±91.79), HTau*^R406W^* (p<0.0001, 456.69±199.30) showed reduced, and APP (p<0.0001, 1016±127.88) showed increased total sleep compared to CC flies. Among the AA flies HTau*^R406W^* (p<0.0001) showed reduced and APP (p<0.0001) showed increased total sleep compared to AA control flies. Day bout numbers (SI. Figure. 4A) increased in APP compared to control in CC flies. In AA flies GFP, HTau and a*β*42 showed increased day bout numbers compared to AA control and HTau*^R406W^*, APP showed reduced day bout numbers compared to CC flies. Night bout numbers (SI. Figure. 4B) increased in phospho HTau compared to Control in CC flies. In AA flies GFP, HTau*^R406W^* showed increased night bout numbers compared to AA control and APP showed reduced night bout numbers compared to CC counterpart. Total bout numbers (SI. Figure. 4C) increased in a*β*42 and APP flies compared to control in CC flies and APP in AA flies showed reduction compared to APP in CC flies. Among the AA flies GFP, HTau, and phospho HTau showed increased total bout numbers compared to AA control flies. Day bout length (SI. Figure. 4D) reduced in all AD flies in CC compared to control except a*β*42 and in AA flies HTau*^R406W^* showed reduced and APP flies showed increased day bout length compared to CC flies. Among the AA flies GFP, HTau, HTau*^R406W^*, and a*β*42 showed significant reduction in day bout length compared to AA control flies. Night bout length (SI. Figure. 4E) reduced in all AD flies compared to control in CC flies and in AA flies GFP shoed reduction and a*β*42, APP showed increased night bout length compared to CC flies. Among the AA flies GFP, HTau, HTau*^R406W^*, and phospho HTau flies showed reduced night bout length compared to AA control flies. Total bout length (SI. Figure. 4F) reduced in all AD flies except a*β*42 in CC flies, in AA flies HTau*^R406W^* showed reduced, and APP showed increased total bout length compared to CC flies. Within AA flies GFP, HTau, and HTau*^R406W^* showed significant reduction in total bout length compared to AA control flies. Day sleep fragmentation (Figure. 4H) increased in HTau*^R406W^*(p<0.02, 0.484±0.22) and APP (p<0.0001, 0.624±0.299) flies in CC condition compared to control (0.291±0.173), and in AA flies APP (p<0.0001, 0.278±0.11) showed reduction compared to CC condition. In AA flies GFP (p<0.0001, 0.482±0.138), HTau*^R406W^*(p<0.003, 0.471±0.232), and a*β*42 (p<0.0001, 0.591±0.323) flies showed increased day sleep fragmentation compared to AA control flies. Night sleep fragmentation (Figure. 4I) increased in a*β*42 (p<0.035, 0673±378) and APP (p<0.049, 0.624±0.239) in CC flies compared to control (0415±0.282) and in AA flies APP (p<0.021, 0.312±0.172) showed reduction compared and GFP (p<0.0001, 0.70±0.23), HTau (p<0.024, 0.504±0.258), HTau*^R406W^* (p<0.0001, 0.874±0.422) showed increased night sleep fragmentation compared to CC flies. Among the AA flies GFP (p<0.001) and HTau*^R406W^* (p<0.0001) flies showed increased night-sleep fragmentation compared to AA control (0.283±0.115). Total sleep fragmentation (Figure. 4J) showed significant increase in HTau*^R406W^* (p<0.021, 1.01±0.37), a*β*42 (p<0.0027, 1.06±0.58) and APP (p<0.0001, 1.31±0.53) in CC flies compared to control (0.66±0.37), whereas in AA flies increase in GFP (p<0.01, 1.16±0.27), HTau*^R406W^* (p<0.02, 1.35±0.62), and decrease APP (p<0.0001, 0.557±0.23) flies compared to CC flies. Among the AA flies GFP (p<0.0001, 1.16±0.27), HTau (p<0.0003, 0.99±0.35), HTau*^R406W^* (p<0.0001, 1.35±0.62) and a*β*42 (p<0.0045, 0.918±0.385) showed increase total sleep fragmentation compared to AA control (0.525±0.165) flies. AD flies found to have disrupted sleep and circadian activity and sleep fragmentation across genotypes. Loss of gut microbiome further exacerbating these disturbances and increasing sleep fragmentation. In APP and a*β*42 flies axenic flies showed differential response indicating the gene-specific interactions of microbiome. Overall gut microbiome influence on sleep/circadian activity suggesting its possible involvement in maintaining the behavioral rhythms during AD pathology.

**Figure 4.**
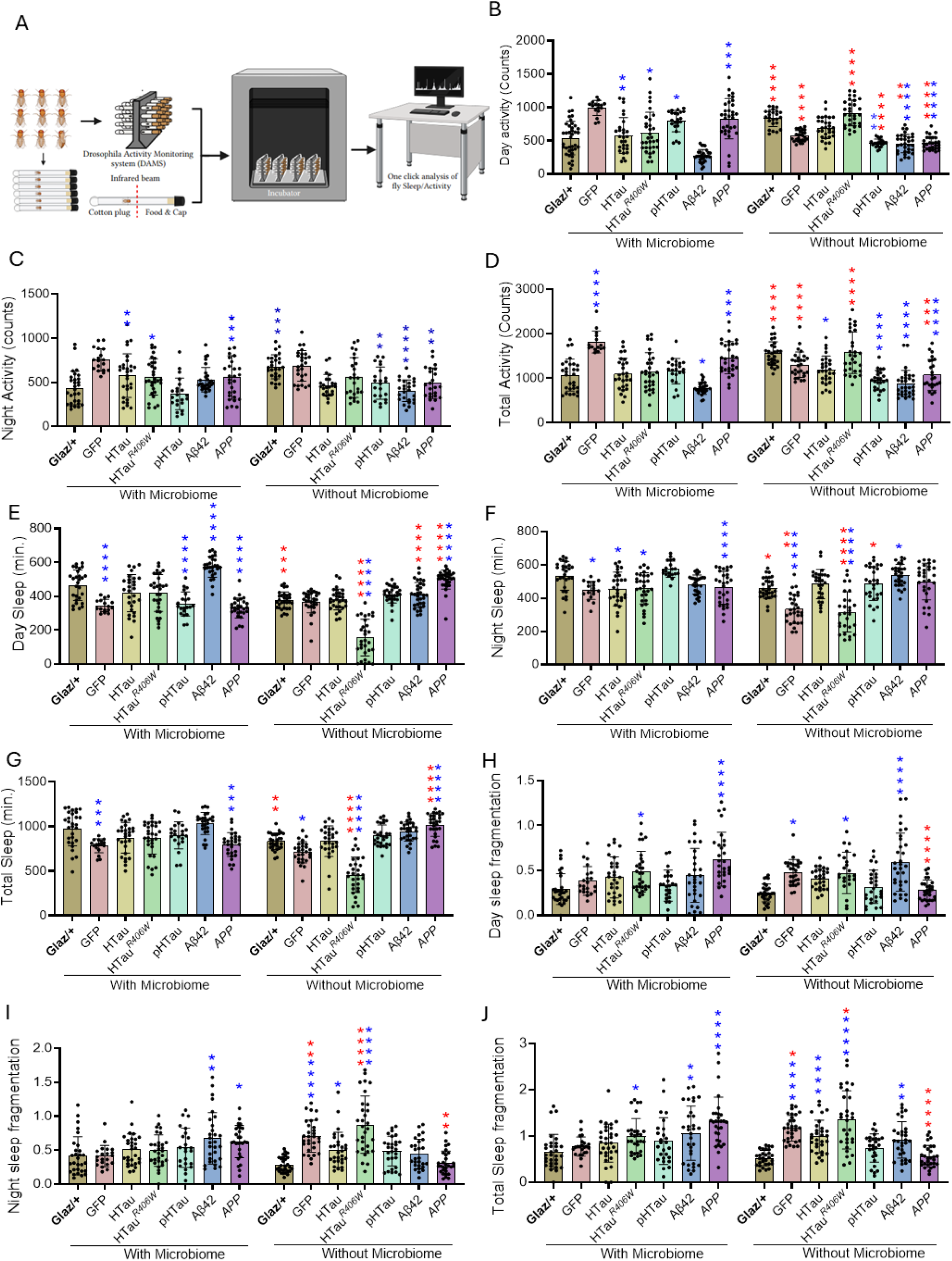
Gut microbiome regulates sleep quality during Alzheimer’s disease. The sleep/activity rhythms of CC and AA flies measure across 24 hours for 3 consecutive days at 3 weeks age. (A). schematic representation of sleep/activity assay using *Drosophila* activity monitoring systems and one click analysis. (B). Day activity increased in HTau (p<0.01), HTau*^R406W^*, phospho HTau (p<0.05), and APP (p<0.001) in CC flies and in AA flies decreased in phospho HTau (p<0.01), a*β*42, and APP (p<0.0001) compared to controls. Compared to CC flies’ day activity increased in *Glaz*/+ and HTau*^R406W^* (p<0.0001) and decreased in GFP, phospho HTau, APP (p<0.0001), and a*β*42 (p<0.01) in AA flies. (C). Night activity increased in (HTau (P<0.01), HTauR*^406W^* (p<0.05), and APP (p<0.001) in CC flies compared to controls. In AA flies *Glaz*/+(p<0.001) showed increased and phospho HTau, APP (p<0.01), a*β*42 (p<0.0001) showed decreased night activity compared to CC flies. (D). Total activity increased in APP (p<0.0001) and decreased in a*β*42 (p<0.05) in CC flies, in AA flies HTau (p<0.01), phospho HTau, a*β*42 and APP (p<0.0001) showed a significant decrease compared to control. Also, total activity increased in *Glaz*/+ and HTau*^R406W^* (p<0.0001) and decreased in GFP (p<0.0001), APP (p<0.001) compared to CC flies. (E). Day sleep decreased in GFP, phospho HTau and APP (p<0.0001) and increased in a*β*42 (p<0.0001) in CC flies, in AA flies decreased in HTau*^R406W^* (p<0.0001) and increased in APP (p<0.0001) compared to control. Compared to CC flies *Glaz*/+, HTau*^R406W^*, a*β*42 showed reduced and in APP (p<0.0001) increased day sleep in AA flies. (F). Night sleep reduced in GFP, HTau, HTau*^R406W^* (p<0.05), and increased in APP (p<0.0001) in CC flies, and in AA flies GFP, HTau*^R406W^* (p<0.0001) showed decreased and a*β*42 (p<0.05) showed increased compared to controls. Compared to CC flies in AA flies *Glaz*/+, phospho HTau (p<0.05), GFP (p<0.01), HTau*^R406W^* (p<0.0001) decreased night activity observed. (G). Total sleep decreased in GFP and APP (p<0.001) in CC flies, in AA flies GFP (p<0.05), HTau*^R406W^* (p<0.0001) and increased in APP (p<0.0001) compared to controls. In AA flies *Glaz*/+ (p<0.01), HTau*^R406W^*(p<0.0001) showed significant reduction and APP (p<0.0001) significant increase compared to CC flies. (H). Day sleep fragmentation increased in HTau*^R406W^* (p<0.05), APP (p<0.0001) in CC flies, in AA flies GFP, HTau*^R406W^* (p<0.05), and a*β*42 (p<0.0001) significantly increased compared to controls and APP (p<0.0001) showed significant reduction compared to CC flies. (I). Night sleep fragmentation increased in a*β*42 (p<0.05) in CC flies, in AA GFP, HTau*^R406W^* (p<0.0001) showed significant increase compared to controls, and APP (p<0.01) compared to CC flies. (J). Total sleep fragmentation increased in HTau*^R406W^* (p<0.05), a*β*42 (p<0.01), And APP (p<0.0001) compared to control, in AA flies GFP, HTau, HTau*^R406W^*(p<0.0001), a*β*42 (p<0.01) showed significant increase compared to control. In APP (p<0.0001) significant reduction in AA flies was observed compared to CC flies. Two-way ANOVA to establish significance, followed by Sidak’s multiple comparisons test to assess the effects of genotype and microbiome condition. n=3, and p-values * <0.05, ** <0.01, *** < 0.001, **** <0.0001.

### 4.5. Depletion of microbiome leads to further compromising AD-linked memory impairment

Based on the immunofluorescence (Figure. 2E) for synapsin and western blot (Figure. 6I) for Synaptotagmin data we understand the possibility of memory impairment in AD flies. We have performed olfaction preference assay with electric shock interruption to quantify the learning behavior in both CC and AA flies. Figure 5A is the schematic representation of memory assay setup. In CC flies (Figure. 5B) before and after training/electric shock *Glaz*/+ (53.57/22.73), GFP (59.46/34.38), HTau (60/33) flies showed reduced preference to same odor after electric shock whereas HTau*^R406W^* (50/53.85), phospho HTau (56.41/54.29), a*β*42 (63.64/60.53) and APP (54.35/54.29) flies showed no change in odor preference/did not learn even after electric shock. This data suggests memory impairment in HTau*^R406W^*, phospho HTau, a*β*42 and APP in CC flies. In AA flies we found a significant memory impairment in all the AD linked genotypes (Figure. 5C) compared to control (60/31.25) and GFP (64/28) flies in HTau (63.64/62.5), HTau*^R406W^* (65/56.25), phospho HTau (60/58.33), a*β*42 (66.67/61), and APP (50/52.38) based on odor choice. We then calculated the olfactory performance index based on the odor choice (Figure. 5D), where negative value represents better performance/no memory impairment and positive values represent severe memory impairments. We represent CC/AA olfactory preference index for *Glaz/+* (–0.47/–0.33), GFP (–0.26/–0.28), HTau (–0.16/0.11), HTau*^R406W^* (0.11/0.21), phospho HTau (0.13/0.21), a*β*42 (0.14/0.45) and APP (0.28/0.11) AD genotypes. We then calculated the avoidance index (Figure. 5E), where higher values represent better memory performance and lesser values represent memory impairment. CC/AA avoidance index for AD files *Glaz/+* (73.52/58.33) GFP (59.6/61.9), HTau (53.3/38.8), HTau*^R406W^* (32.4/30), phospho HTau (37.8/34.7), a*β*42 (37.12/22.72) and APP (31.57/38.88). The olfactory preference assay showed clear learning/memory phenotypes in some genotypes in CC flies, and the axenic flies showed exacerbated memory loss in all AD models. The performance index and avoidance index further confirm the possible impact of loss of gut microbiome further worsening the cognitive decline in AD models.

**Figure 5.**
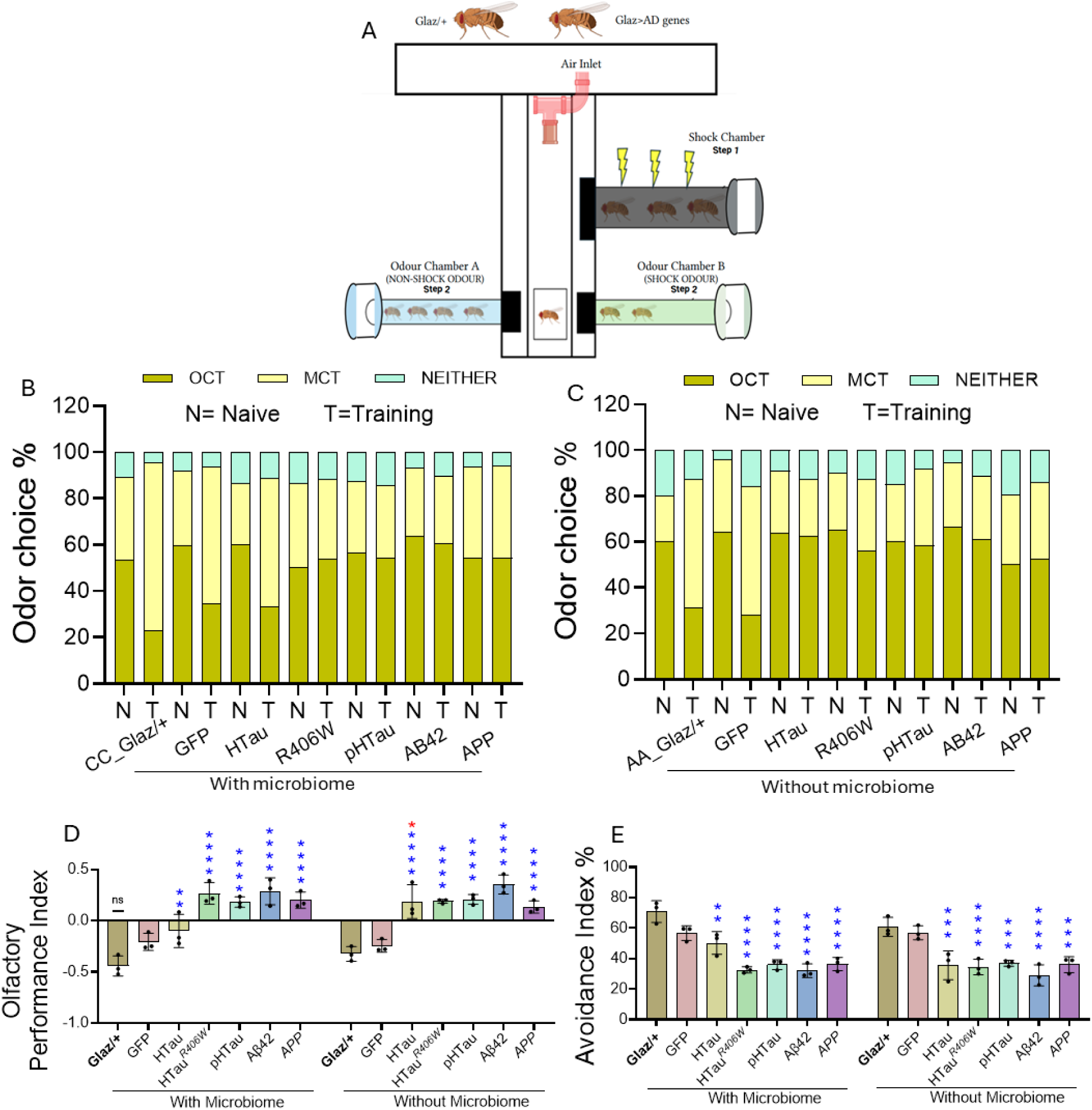
Depletion of microbiome worsening the memory impairment linked with AD. Gut microbiome influence on memory/cognitive performance in AD flies tested using OCT (3-Octanol) and MCT (4-Methylcyclohexanol) odors and under the influence of electric shock (110 Volts) at 3 weeks age. (A). Schematic representation of memory assay for *Drosophila,* (B). Comparison of odor choice% between Naïve (N)-before shock and Training (T) after shock in conventional control (CC)/with microbiome AD and control flies. If flies still prefer shock odor, it implies memory impairment. (C). Odor choice % in axenic antibiotic (AA)/without microbiome AD and control flies, (D). Olfactory performance index % showed a significant increase in HTau (p<0.01), HTau, HTau*^R406W^*, phospho HTau, a*β*42 and APP (p<0.0001) in CC flies, in AA all the AD flies showed significant increase (p<0.0001) compared to control, and HTau(p<0.05) showed significant increase compared to CC flies. (E). Avoidance index % significantly reduced in CC (p<0.0001), in AA (p<0.0001) AD flies compared to controls. memory assay panel B, and C chi-square test for panel D, and E Two-Way ANOVA with Sidak’s multiple comparison test was used. n=3, and p-values * <0.05, ** <0.01, *** < 0.001, **** <0.0001.

**Figure 6.**
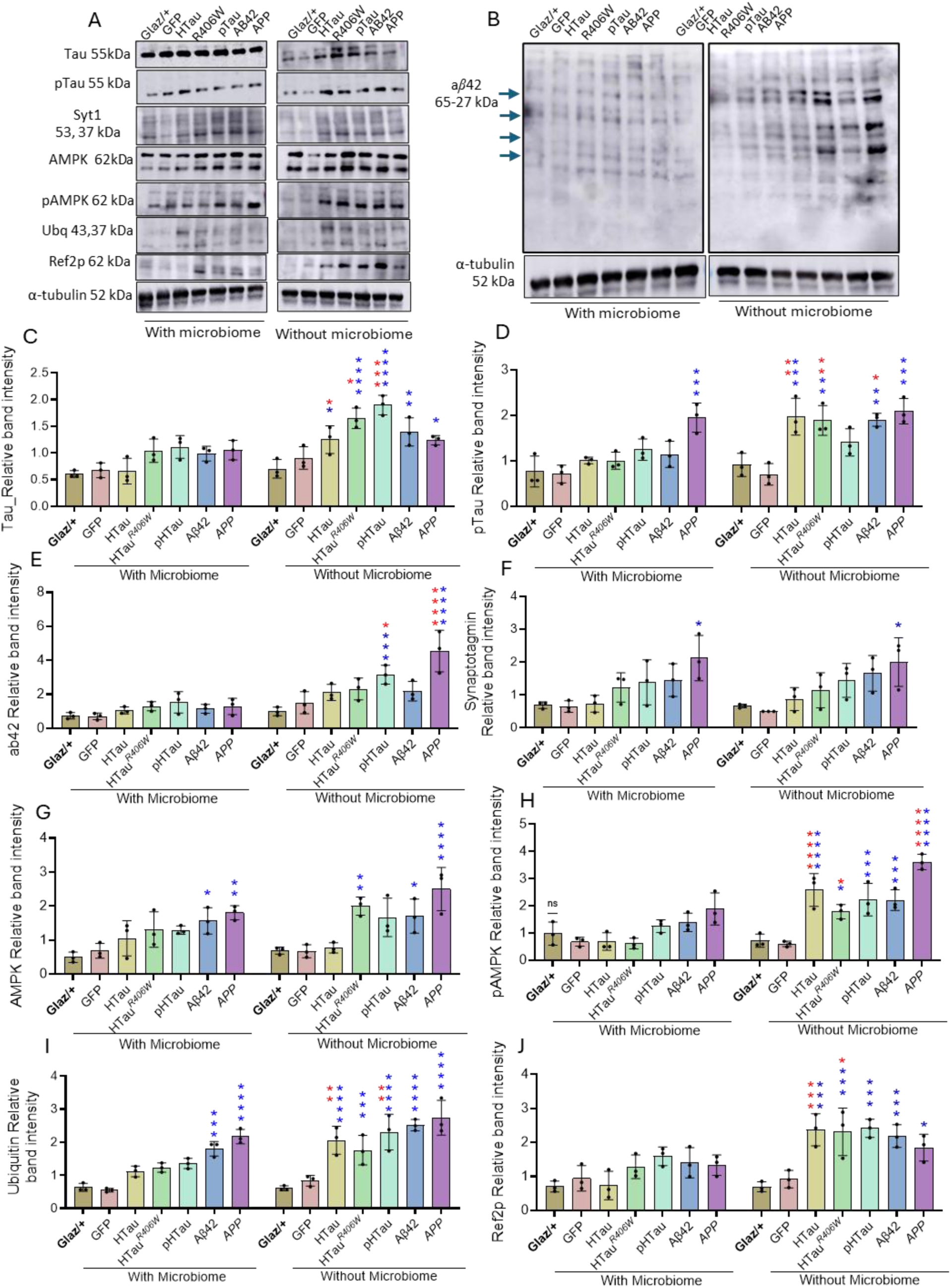
Depletion of microbiome resulted in upregulation of a*β*42 and phospho Tau through impaired autophagy mediated by activation of AMPK. Western blot analysis to quantify the abundance of targeted proteins in the head (C). Tau band intensity shows no significant difference in CC flies, in AA flies HTau, APP (p<0.05), HTau*^R406W^*, phospho HTau (p<0.0001) and a*β*42 (p<0.01) compared to controls. In AA flies compared to CC, HTau, HTau*^R406W^* (p<0.01), and phospho HTau (p<0.001) showed significant increase. (D). phospho Tau increased in APP (p<0.001) in CC flies, in AA flies HTau, APP (p<0.001), and HTau*^R406W^*, a*β*42 (p<0.01) showed significant increase compared to controls. AA, HTau and HTau*^R406W^* (p<0.01) showed significant increase compared to CC flies. (E). a*β*42 band intensity increased in phospho HTau (p<0.001) and APP (p<0.0001) in AA flies compared to control and phospho HTau (p<0.05), APP (p<0.0001) compared to CC flies. (F). Syt1 increased in APP (p<0.05) in both CC and AA flies with no significant differences. (G). AMPK band intensity increased in a*β*42 (p<0.05) and APP (p<0.01) in CC, and in AA HTau*^R406W^*, a*β*42 (p<0.05), and APP ((p<0.0001) compared to controls. (H). phospho AMPK, has no significant change in CC flies, in AA HTau, phospho HTau, a*β*42, APP (p<0.0001), HTau*^R406W^* (p<0.05) compared to controls. In AA flies compared to CC flies HTau (p<0.0001), and HTau*^R406W^* (p<0.05) showed significant increase. (I). Ubiquitinated proteins increased in a*β*42 (p<0.001) and APP (p<0.0001) in CC flies, in AA flies HTau, phospho HTau, a*β*42, APP (p<0.0001), HTau*^R406W^* (p<0.0015) compared to controls. In AA flies compared to CC flies HTau and phospho HTau (p<0.01) showed significant increase. (J). Ref2p (p62 homologue of *Drosophila melanogaster*) has no significant difference in CC flies, but in AA flies HTau, HTau*^R406W^*, phospho HTau, a*β*42 (p<0.001), and APP (p<0.05) showed significant increase in band intensity compared to controls. Compared to CC flies AA, HTau (p<0.001), HTau*^R406W^*(p<0.01) showed significant increase. The representative blots (A-B) were shown along with internal control alpha tubulin. Two-way ANOVA to establish significance, followed by Sidak’s multiple comparisons test to assess the effects of genotype and microbiome condition. n=3, and p-values * <0.05, ** <0.01, *** < 0.001, **** <0.0001.

We then also performed geotaxis analysis in male (SI. Figure. 5A-F) and female (SI. Figure. 6A-F) flies of CC and AA condition. Line graph represents climbing distance of each genotype from 0 to 9 seconds. We have observed a significant reduction in locomotory performance in AA, GFP, HTau*^R406W^*, and phospho HTau flies compared to CC GFP, HTau*^R406W^*, and phospho HTau flies. No significant difference in HTau AA flies compared to CC counterparts. But a*β*42 and APP showed a significant increase in AA flies compared to CC flies in both males and females. In females HTau and HTau*^R406W^* showed significant change in locomotory performances in AA flies compared to CC flies. These data suggested that microbiome depletion differentially influencing the climbing behavior (geotaxis responses) in AD transgenic flies, suggest potential genotype–specific interactions of host and microbiome could influence geotaxis performance.

### 4.6. Depletion of microbiome leads to enhanced pAMPK, compromised autophagy, and increased p-tau and a*β*-42 accumulation suggesting defective gut-brain communication

To understand how microbiome absence impacts the energy status and metabolism, we have checked the AMPK (Figure. 6G) protein band intensity in AA and CC flies. In CC a*β*42 (p<0.02, 1.56±0.38) and APP (p<0.003, 1.81±0.21) flies showed increase AMPK band intensity than control (0.49±0.15). In AA fliesHTau*^R406W^* (p<0.003, 1.99±0.26), a*β*42 (p<0.04, 1.71±0.50) and APP (p<0.0001, 2.49±063) showed significant increase compared to AA control (0.68±0.10). Active form of AMPK i.e., phospho AMPK (Figure.6H) did not show any significant increase in CC flies compared to control, but in AA flies HTau (p<0.003, 2.58±0.61), HTau*^R406W^*(p<0.032, 1.78±0.26), phospho HTau (p<0.0007, 2.23±059), a*β*42 (p<0.0007, 2.21±0.37) and APP (p<0.0001, 3.61±0.28) flies showed a significant increase compared to AA control (0.72±0.24) flies. In addition, we observed a significant increase in pAMPK in HTau (p<0.0001, 0.68±0.32), HTau*^R406W^* (p<0.012, 0.61±0.19) and APP (p<0.0001, 3.61±0.28) compared to CC flies, which suggests the possible deprivation of energy status during axenic condition, which may be due to poor absorption of food from the gut^50,51^. Since pAMPK can promote the autophagy mechanism, which is essential for reducing the protein aggregates building up in AD condition by breaking them into precursor molecules^52,53^. So, we have checked the ubiquitinated protein (Figure. 6I) band intensity in flies. In CC flies a*β*42 (p<0.0007, 1.81±0.22) and APP (p<0.0001, 2.17±0.21) showed a significant increase in ubiquitinated protein band intensity compared to control (0.63±0.12). In AA flies HTau (p<0.0093, 2.05±0.42) and phospho HTau (p<0.0083, 2.30±0.53) flies showed significant increase compared to CC flies. In HTau (p<0.0001, 2.05±0.42), HTau*^R406W^* (p<0.0009, 1.75±0.44), phospho HTau (p<0.0001, 2.30±0.53), a*β*42 (p<0.0001, 2.51±0.6) and APP (p<0.0001, 2.73±0.52) flies have a significant increase compared to AA control (0.61±0.07) flies. We then observed an autophagy receptor Ref2p (p62) which directs damaged proteins to autophagosome^54,55^. In our study we did not observe any significant increase in CC flies compared to control (Figure. 6J), in AA condition HTau (p<0.0002, 2.30±0.47) and HTau*^R406W^* (p<0.041, 2.31±0.46) flies showed an increased accumulation of Ref2p compared to CC flies (0.71±0.15). Whereas HTau (p<0.0003, 2.30±0.47), HTau*^R406W^* (p<0.0002, 2.31±0.69), phospho HTau (p<0.0001, 2.41±0.26, a*β*42 (p<0.0007, 2.18±0.33) and APP (p<0.014, 1.84±0.39) flies showed significant increase in Ref2p compared to AA control (0.69±0.14) flies. Our results suggest that probability of greater involvement of gut microbiomes in proteostasis in AD flies.

Since gut microbiome significantly impacting lipid accumulation, neuroinflammation, sleep quality and impaired proteostasis, we further focused on AD associated protein expression patterns in both CC and AA flies (Figure. 6). Tau protein band intensity (Figure. 6C) in CC flies did not show any significant difference in AD flies compared to CC control flies. In AA flies HTau (p<0.032, 1.24±0.26), HTau*^R406W^* (p<0.0001, 1.64±0.19), phospho HTau (p<0.0001, 1.891±08) and a*β*42 (p<0.0026, 1.39±0.26) and APP (p<0.037, 1.29±0.86) flies showed significant increase compared to AA control (0.72±0.17) flies. In AA flies HTau (p<0.015), HTau*^R406W^*(p<0.011), phospho HTau (p<0.0005) showed a significant increase in Tau protein band intensity compared to CC HTau (0.65±0.24), HTau*^R406W^* (1.032±0.21), phospho HTau (1.11±0.21) flies. We then checked the phospho HTau protein levels (Figure. 6D), in CC flies APP (p<0.0002, 1.95±0.32) showed a significant increase compared to CC control (0.76±0.34) flies. In AA flies HTau (p<0.0009, 1.97±0.41), HTau*^R406W^* (p<0.0024, 1.88±0.32), a*β*42 (p<0.002, 1.99±0.14) and APP (p<0.0002, 2.09±0.28) showed a significant increase in phospho HTau band intensity compared to AA control (0.91±0.25) flies. In HTau (p<0.0029), HTau*^R406W^*(p<0.0074) and a*β*42 (p<0.033) flies we found a significant increase in phospho HTau compared to CC, HTau (1.00±0.07), HTau*^R406W^* (100±0.18) and a*β*42 (1.01±0.28). We further checked the a*β*42 protein band intensity (Figure. 6E), in CC condition we did not observe a significant increase in a*β*42 protein band intensity in AD flies compared to control (0.73±0.18). In AA flies phospho HTau (p<0.0007, 3.15±0.56) and APP (p<0.0001, 4.54±1.22) showed a significant increase compared to AA control (0.99±0.25) flies. Further we found a significant increase in phospho HTau (p<0.0007, 3.15±0.56) and APP (p<0.0001, 4.54±1.22) in AA flies compared to CC phospho HTau (1.52±0.62) and APP (1.28±0.48) flies. We have checked the synaptotagmin1 (Syt1) protein band intensity (Figure. 6F), a presynaptic protein essential for release of synaptic vesicles into synaptic cleft^56^. We have observed a significant increase in band intensity of Syt1 in APP (p<0.014, 2.11±0.68) compared to control (0.68±0.10) in CC and in AA flies (p<0.027, 1.99±0.73) compared to control (0.65±0.056). As we have observed increase in tau, phospho HTau and a*β*42 protein band intensity in many AD fly models and increased pAMPK, a positive regulator of the autophagy suggests possibility of impaired autophagy in AD flies especially in Axenic condition. Elevated Tau, phospho HTau, a*β*42 protein levels along with ubiquitination and Ref2p accumulation suggests greater risk of protein aggregation in axenic flies. Although elevation of AMPK and pAMPK could suggest energy stress pathways, increased levels of protein aggregation suggest possible disconnection in downstream autophagy clearance. To further confirm we have performed Atg8a/LC3-I and Atg8b/LC3-II forms which involved recruiting proteins and forming autophagosome to complete the protein degradation (SI Figure 6A). We found a significant increase in LC3-I in AD samples during AA condition and negligible quantities of LC3-II protein band intensity. At the same time CC flies LC3-II band is apparently visible. This highlights gut microbiome’s modulation of energy metabolism and autophagy mechanism essential for clearance of protein aggregates. This interconnection observed between gut microbiome depletion and impaired protein homeostasis emphasizes significance of gut microbiome in buffering neurodegenerative stress during AD pathology.

## 5. Discussion

Alzheimer’s disease has been discussed as a central nervous system disorder driven by amyloid beta plaques and hyperphosphorylated tau tangles^57,58^. However large amount of evidence supports that AD is influenced by systemic processes, including peripheral immune responses, metabolic regulation and gut to brain communication^7,59,60^. Our study provides experimental evidence to support this idea, where depletion of gut microbiome in humanized *Drosophila* model of AD aggravates disease pathology at molecular, cellular and behavioral levels. By comparing AD flies growing under axenic antibiotic (AA) and conventional control (CC) conditions our data supports that the microbiome can function as an active modulator in controlling the neurodegenerative stress. Our study showed altered microbiome population in the gut of AD transgenic flies, where *Drosophila* abundant microbiome *Lactobacillus* and *Acetobacter* showed marked differences in the AD transgenic flies. This observation aligns with recent human and murine studies showing that dysbiosis associated with AD varies depending on disease stage, host genetics, and metabolic state. For example, alterations in *Lactobacillus, Bifidobacterium*, and pro-inflammatory taxa such as *Escherichia/Shigella* have been linked to increased peripheral inflammation and cognitive decline^61,62^. Since *Drosophila* AD model recapitulates microbial shift (Figure 1; S.I. 1) like previous studies^61,62^, our data supports the relevance of flies as an idle model for studying gut-brain communication during AD pathology.

One of the fundamental observations we found in this study was that removal of microbiota leads to significant upregulation of proinflammatory cytokines and their downstream JAK/STAT signaling (Figure.3; S.I. 2). In AD flies, we observed upregulation of *Upd-Dome-Hop-Stat92e* axis. This pathway is functionally analogous to mammalian IL-6/JAK/STAT signaling, which is strongly implicated in AD-associated neuroinflammation and microglial activation^63^. This pathway involved activation of microglia^64^ and even synaptic dysfunction in AD^63^. Our findings suggest the possibility of potential link between microbiome depletion and inflammatory signaling activation, but further studies are required to establish the mechanistic relationship and intermediates involved. Studies in mammalian models showed microbial metabolites specifically short chain fatty acids (SCFA) such as acetate, butyrate suppresses inflammatory responses driven by NF-kB and STAT signaling^64,65^. Which supports dysbiosis/depletion of gut microbiome could contribute to reduction of SCFA. Lake of microbial derived metabolites further exacerbate the inflammatory responses by reactive gliosis and reduces the clarence of protein aggregates^64,66^ which further accelerates neuronal cell death. Our data showed a simultaneous upregulation of apoptotic mediators such as *Hid* and *Reaper* under axenic conditions in both head and body regions (Figure. 3; S.I. 2) support the idea that upregulation of inflammation directly impacts cell death pathways^67,68^. These data suggests that microbiomes maintain an intricate balance between immune system, neurosignaling and apoptotic stress whereas its depletion intensifies AD pathology. We believe our preliminary findings align with mammalian inflammatory signaling activation, but further studies are required to confirm this link to microbial depletion.

Growing evidence support connection between AD and metabolic disorders, with strong association to insulin metabolism, lipid dysregulation and mitochondrial dysfunction^69–71^. In AD, a consistent pathological signature includes increased phosphorylated AMPK (p-AMPK), accumulation of p62 (Ref2P), and elevated levels of ubiquitinated proteins, reflecting impaired proteostasis. AMPK activation indicates chronic neuronal energy stress and promotes autophagy initiation^72^, while p62, an adaptor that binds ubiquitinated cargo, is normally degraded during autophagy and therefore accumulates when autophagic flux is defective^73^. The concurrent increase in ubiquitinated protein aggregates, a hallmark of AD pathology found in neurofibrillary tangles and dystrophic neurites, further supports a failure in protein clearance systems^74^. Our study further extends this view by showing increased lipid droplet accumulation, Ref2p/p62, and LC3-I/LC3-II ratio when gut microbiome is depleted in AD transgenic flies. Increased lipid droplets in the head and brain regions further supports the possible dysregulated lipid metabolism or altered lipid homeostasis^75,76^, where gut microbiome derived acetate is known to play a significant role^77,78^. It also correlates with results observed in both murine models and human AD patients^79^. Interestingly, axenic AD flies showed increased AMPK phosphorylation, a master regulator that checks cellular energy balance^52^. Collectively, the increase in p-AMPK, ubiquitinated proteins, Ref2p, and LC3-I with reduced LC3-II suggests a possible impairment in autophagic flux and defective protein degradation^52,55,80,81^. Based on our study on axenic flies we believe that absence of gut microbiome might disrupt the nutrient absorption in the gut and availability of microbial metabolites^50^. Hence, energy stress condition promoting AMPK activation but failing to accomplish autophagy clearance of aggregated proteins^82^. Another possibility lies within the AMPK dependent activation of autophagy mechanism, AMPK regulates ULK1 under glucose starvation and amino acid starvation differently^52^. Our findings further supported by recent studies emphasizing the role of microbiome-derived metabolites involved in enhancing the lysosomal and autophagy functions^83–85^. Together these findings indicate that autophagy is activated but not efficiently completed, leading to accumulation of ubiquitinated proteins and contributing to neurodegeneration^86,87^. Based on these insights, we believe that the gut microbiome could help to mitigate AD pathology by supporting both lipid metabolism and autophagic mechanisms. Depletion of gut microbiome could potentially suppress these protective inputs, that leads to impaired proteostasis neurofibrillary tangles formation, dystrophic neuritis and amyloid accumulation^88^.

It is well known that AD patients were prone to show sleep fragmentation, changes in activity cycle, abnormal eating patterns^89,90^. In our study we have observed a significant sleep/circadian disruption in axenic AD fly models compared to CC flies (Figure. 4; S.I. 3). Which suggests a strong correlation between gut microbiome and behavioral patterns. Apart from that, gut microbiome itself exhibits diurnal oscillations in microbial composition, metabolite production, that could interact with host internal circadian clock proteins^91^. This relationship strongly suggests the possible missing of an element particularly due to depletion of microbiota that perhaps synchronizes host circadian machinery. One convincing mechanism that support our interpretation is that microbial metabolites could act as a zeitgebers, entraining peripheral clocks through AMPK, sirtuins and nuclear hormone receptors^92–95^. Loss of these cues may aggravate the circadian misalignment that was already established by tauopathy and amyloid pathology and aggravating metabolic and cognitive dysfunctions^96^. Therefore, our results support the concept of bidirectional relationship of gut-brain axis^7^. As AD pathology disrupts circadian rhythms, which in turn reshape the gut microbiome^97^, and disruption of gut microbiome further amplifies cognitive decline and circadian misalignment^98^. This never-ending loop could partly explain the progressive worsening of behavioral changes observed in AD patients.

One of the significant pathological associations of cognitive decline in AD condition is synaptic loss^99^. We have observed a significant reduction in the synapsin in the axenic flies^100^ alongside impaired olfactory preference and shock avoidance memory (Figure. 5). It could be due to limited support available from gut microbiota during microbiome depletion in axenic condition^101^ and destabilize synaptic communication by limiting metabolic and trophic support^102–105^. Several direct and indirect evidence support this interpretation, since neurotransmitter precursors such as tryptophan, dopamine, and gamma amino butyric acid found to be influenced by microbial metabolites^106^. SCFA could promote the synaptic protein expression and spine maturation^107^. Dysbiosis could lead to decreased synaptic markers and even memory performance^108^. All these data support our understanding of microbiome influence on synaptic stability. In addition, we have observed all AD flies exhibited cognitive impairment during axenic condition, weighing on microbiome involvement is critical across all the tested AD genotypes. This highlights microbiome modulation could serve as a common therapeutic approach for AD associated memory impairments during Alzheimer’s disease pathology.

An important conceptual insight emerging from this study is that microbiome depletion does not appear to act through a single pathway but rather influences a network of interconnected immune, metabolic, and neuronal processes. Within this context, the gut microbiome may contribute to maintaining homeostasis across these systems. Its depletion is associated with a shift toward a more vulnerable state, characterized by increased inflammatory signaling, altered metabolic balance, impaired proteostasis, and behavioral changes. These effects are unlikely to be independent and may instead interact in a reinforcing manner, potentially contributing to disease progression. This interpretation is broadly consistent with recent cross-species studies^7,109^ suggesting that microbiome perturbations can modulate AD-related phenotypes through coordinated immune–metabolic interactions.

## Limitations and future directions

Although the current study provides new insights into microbiome-related modulation of glial-mediated AD pathology, few limitations must be acknowledged when assessing these results. Though *Drosophila* offers effective genetic resources to analyze glial–microbiome interactions, simplified immune system and microbiome composition might not completely reflect mammalian AD pathology. Our data demonstrates robust connections between microbiome health, circadian disruption, and proteostasis imbalance, yet the exact molecular intermediates that connect microbial signals to glial inflammatory pathways are still to be determined. Future research combining metabolomic profiling and mammalian models will be crucial to extend these results.

## Conclusion

Current manuscript discusses how microbiome depletion exacerbates glial-driven neuroinflammation, sleep/circadian disruption and proteostasis imbalance in AD model of Drosophila, which mainly highlights gut microbiome as an important factor contributing to neurodegenerative phenotype. Our study provides mechanistic preclinical framework suggesting that metabolic homeostasis and glial resilience are highly influenced by microbial signals during AD progression. As our model system enables precise genetic manipulation of gut-brain interactions, future studies using mammalian systems help to confirm the translational relevance of these pathways. We believe our study support the concept of gut microbiome-glial interaction contribute to Alzheimer’s disease pathology and endorse a new direction for future studies in the context of neurodegenerative research.

## Funding/Acknowledgement

This work was supported by National Institutes of Health (NIH) grants AG065992, AGO68550 and RF1NS133378 to G.C.M This is partially supported by UAB startup funds 3123227, 3123226 and UAB Nathan Shock Center P30 AG050886 (USA) to G.C.M.

## Author contribution

Under GCM guidelines, KM designed the experiments, KM performed most of the experiments, GS performed sleep/activity, JW performed immunofluorescence experiments. KM analyzed all the experimental data and wrote the draft of manuscript. GCM edited and revised the figures and manuscript.

## Data availability

The authors confirm that data supporting findings of the current study provided in the manuscript, supplementary material and resource data file.

## Supplementary materials

Supplementary materials for this manuscript can be accessed online.

## Supplementary information

**SI. Figure 1.**
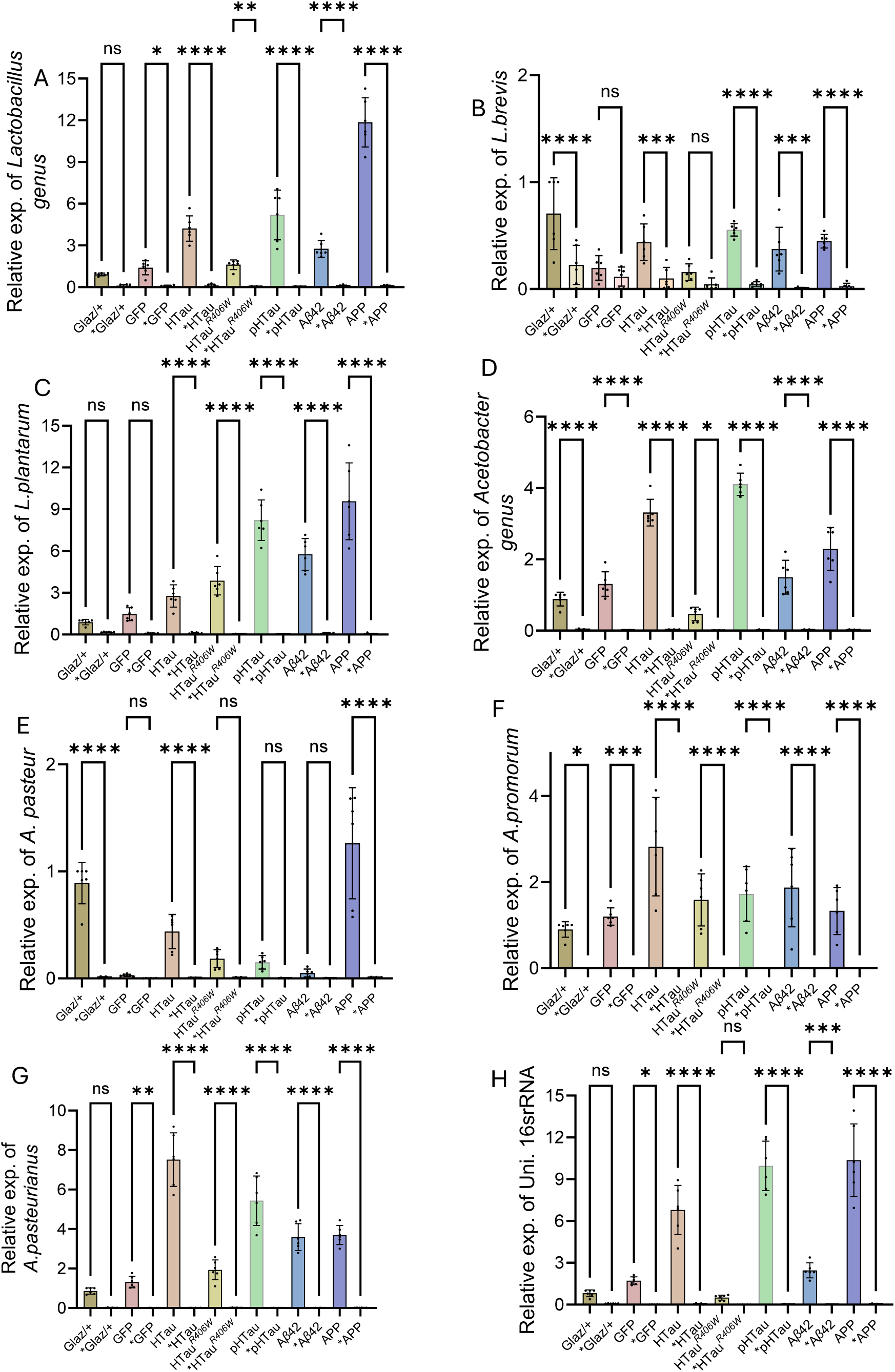
Represents the comparison of most abundant microbial population in fly gut between conventional control (CC) and axenic antibiotic (AA) flies carrying AD transgenes expressed under *Glaz-Gal4* glial cell specific driver at 3 weeks age. (A). *Lactobacillus* genus AA flies showed a significant reduction in HTau, pTau, a*β*42, APP (p<0.0001) and GFP, HTau*^R406W^* (p<0.01) compared to CC flies. (B). *L. brevis* showed significant decrease in AA flies (p<0.0001) compared to CC flies in *Glaz*/+, HTau, pTau, a*β*42, and APP flies. (C). *L. plantarum* Significant reduction in GFP, HTau (p<0.001), HTau*^R406W^*, pTau, a*β*42, APP (p<0.0001) in AA flies compared to CC. (D). *Acetobacter* in AA flies showed a significant reduction in *Glaz*/+, GFP, HTau, pTau, a*β*42, APP (p<0.0001) and HTau*^R406W^* (p<0.01) compared to CC flies. (E). *A. pasteur* in *Glaz*/+, HTau, and APP flies in AA showed significant decrease compared to CC flies. *A. promorum Glaz*/+, GFP (p<0.01) and HTau, HTau*^R406W^*, pTau, a*β*42, APP (p<0.0001) in AA flies decreased significantly compared to CC flies. (G). *A. pasteurianus* showed significant reduction in GFP (p<0.01), HTau, HTau*^R406W^*, pTau, a*β*42, APP (p<0.0001) in AA flies compared to CC flies. (H). 16 rRNA quantification showed significant reduction in GFP (p<0.01), HTau, pTau, a*β*42, APP (p<0.0001) in AA flies compared to CC flies. n=3, p-values * <0.05, ** <0.01, *** < 0.001, **** <0.0001.

**SI. Figure 2.**
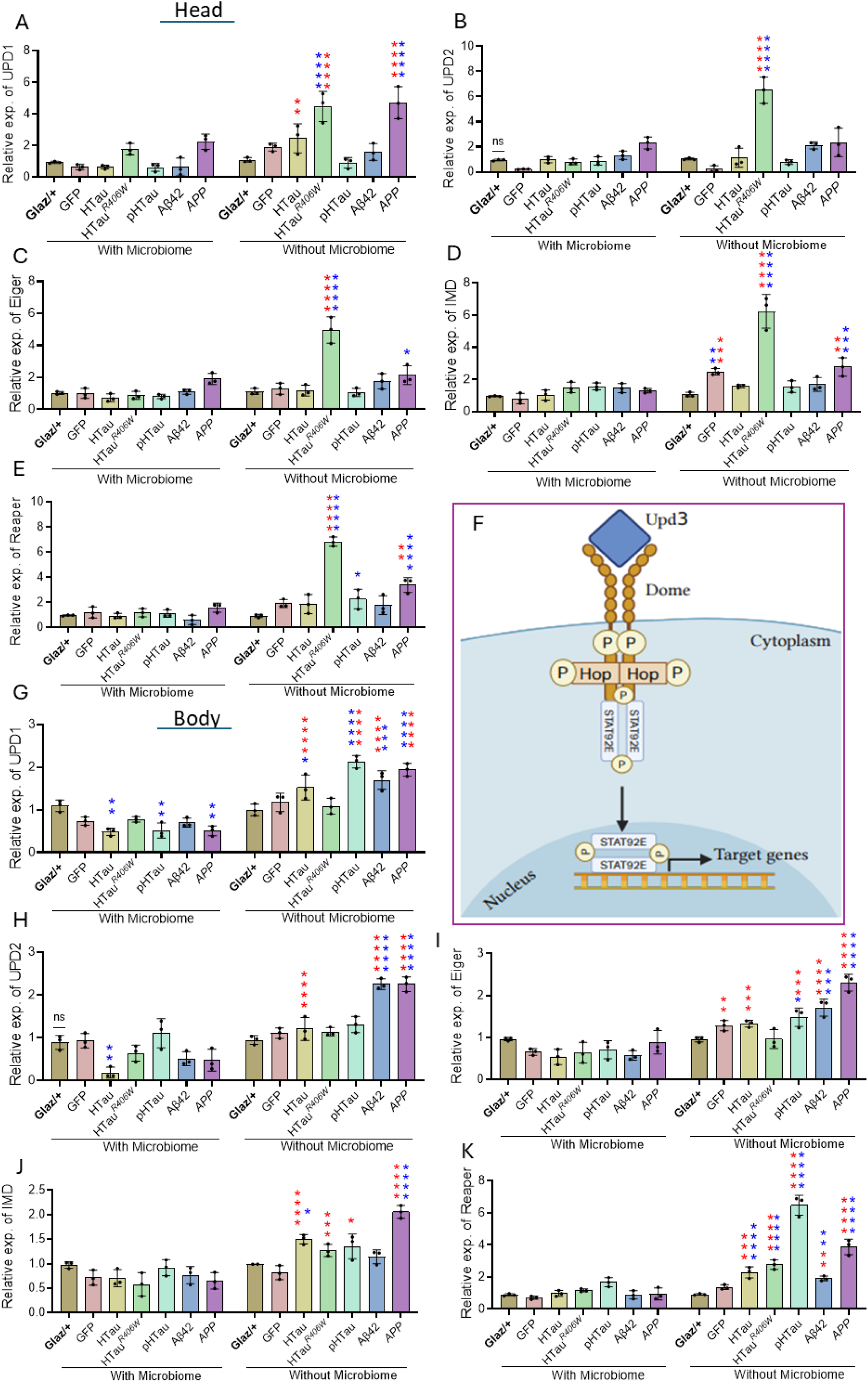
Represents cytokine (*Upd*1, and 2), *Eiger* (*TNFα* homolog) its downstream target *Imd* and cell death marker *Reaper* expression in head and body regions. (A). *Upd*1 expression have no significant difference in CC flies, in AA HTau*^R406W^*, and APP (p<0.0001) showed a significant difference compared to controls. HTau (p<0.01), HTau*^R406W^*, and APP (p<0.0001) showed significant increase compared to CC flies. (B). *Upd*2 expression found to be increased only in HTau*^R406W^* (p<0.0001) compared to control and CC counter part in AA flies. (C). *Eiger,* showed increased expression in HTau*^R406W^* (p<0.0001), and APP (p<0.1) compared to control in AA flies and HTau*^R406W^* (p<0.0001) compared to CC flies. (D). *Imd* increase in GFP (p<0.01), HTau*^R406W^* (p<0.0001), GFP (p<0.001), HTau*^R406W^* (p<0.0001), and APP (p<0.01) compared CC flies. (E). *Reaper* gene expression increased in AA HTau*^R406W^*, APP (p<0.0001), and pTau (p<0.05) compared controls and HTau*^R406W^* (p<0.0001), APP (p<0.01) compared to CC flies in head region. (F). Schematic representation of inflammatory signaling through *Upd 3-Stat92e* pathway. (G). *Upd* 1 expression decreased in HTau, pTau and APP (p<0.01) in CC flies, and HTau (p<0.05), pTau, APP (p<0.0001) and a*β*42 (p<0.001) in AA flies compared to controls. In AA flies HTau, pTau, a*β*42 and APP (p<0.0001) showed significant increase compared to CC flies. (H). *Upd 2* showed significant decrease in HTau (p<0.01) in CC flies, a*β*42 and APP (p<0.0001) in AA flies compared to controls. In AA flies HTau, a*β*42 and APP (p<0.0001) showed a significant increase compared to CC flies.(I). *Eiger* expression increased in pTau (p<0.05), a*β*42 (p<0.001) and APP (p<0.0001) compared to control in AA flies, and Compared to CC flies GFP (p<0.01), HTau, pTau (p<0.001), a*β*42 and APP (p<0.0001) increased significantly in AA flies. (J). Imd upregulated in AA HTau (p<0.05) and APP (p<0.0001) compared to control, also HTau, APP (p<0.0001), HTau*^R406W^* (p<0.001), and pTau (p.0.05) showed significant increase in AA flies compared to CC flies. (K). *Reaper* significantly upregulated in AA, HTau, HTau*^R406W^*, pTau, APP (p<0.0001) and a*β*42 (p<0.01) compared to controls. Also, compared to CC flies HTau (p<0.001), HTau*^R406W^*, pTau, APP (p<0.0001) and a*β*42 (p<0.01) increased significantly in AA flies. n=3, p-values * <0.05, ** <0.01, *** < 0.001, **** <0.0001.

**SI. Figure 3.**
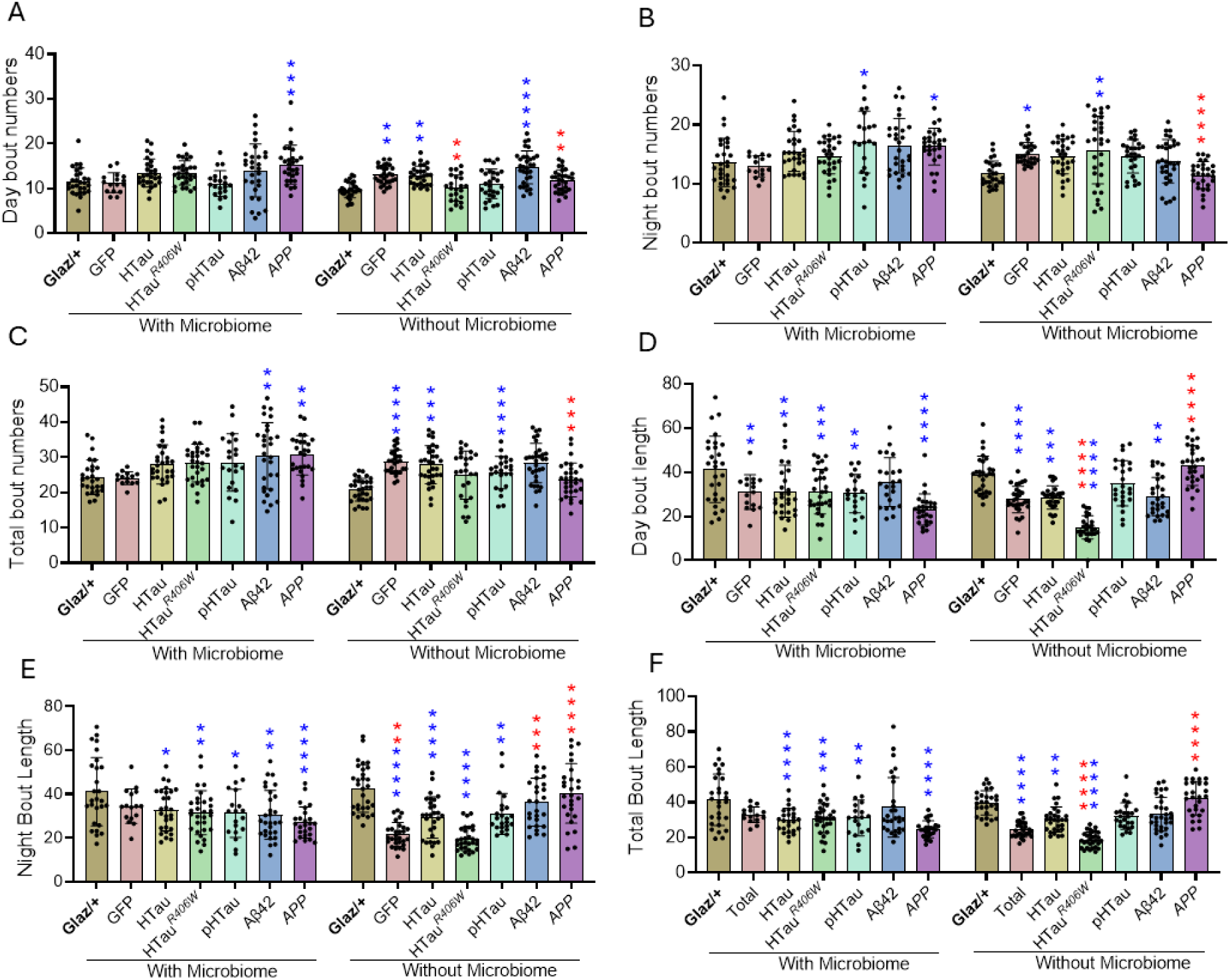
Represents the bout numbers, and bout length. in CC and AA flies at 3 weeks age. (A). Day sleep bout numbers increased in APP flies compared to control in CC, and in AA flies GFP, HTau (p<0.01) and a*β*42 (p<0.0001) compared to controls, (B). Night sleep bout numbers increased in pTau and APP (p<0.05) in CC flies, In AA flies GFP (p<0.05), HTau*^R406W^* (p<0.01) compared to controls. In APP of AA flies significant increase observed (p<0.0001) compared to CC flies. (C). Total sleep bout numbers increase in a*β*42, APP (p<0.0001) of CC flies, in AA flies GFP, pTau (p<0.0001) and HTau (p<0.001) compared to controls. In APP of AA flies significant increase observed (p<0.001) compared to CC flies. (D) Day sleep bout length decreased in GFP, HTau, and pTau (p<0.01), HTau*^R406W^*(p<0.001) and APP (p<0.0001) compared to controls. In AA flies GFP, HTau*^R406W^* (p<0.0001), HTau, a*β*42 (p<0.01) decreased significantly compared to control. In AA flies HTau*^R406W^* (p<0.0001) significantly decreased, and APP (p<0.0001) significantly increased compared to CC flies. (E). Night sleep bout length decreased in HTau, pTau (p<0.05), HTau*^R406W^*, a*β*42 (p<0.01) and APP (p<0.0001) in CC flies compared to control. In AA flies GFP, HTau, HTau*^R406W^* (p<0.0001), and pTau (p<0.01) decreased compared to controls. In AA GFP (p<0.01), decreased and a*β*42 (p<0.001), APP (p<0.0001) significantly increased compared to CC flies. (F). Total sleep bout length Decreased in HTau, APP (p<0.0001), HTau*^R406W^* (p<0.001), and pTau (p<0.01) compared to control I n CC flies. In AA flies GFP, HTau*^R406W^* (p<0.0001), and HTau (p<0.01) decreased significantly compared to control in AA flies. In HTau*^R406W^* (p<0.001) decreased significantly and in APP (p<0.0001) increased significantly compared to CC flies. n=3, p-values * <0.05, ** <0.01, *** < 0.001, **** <0.0001.

**SI. Figure 4.**
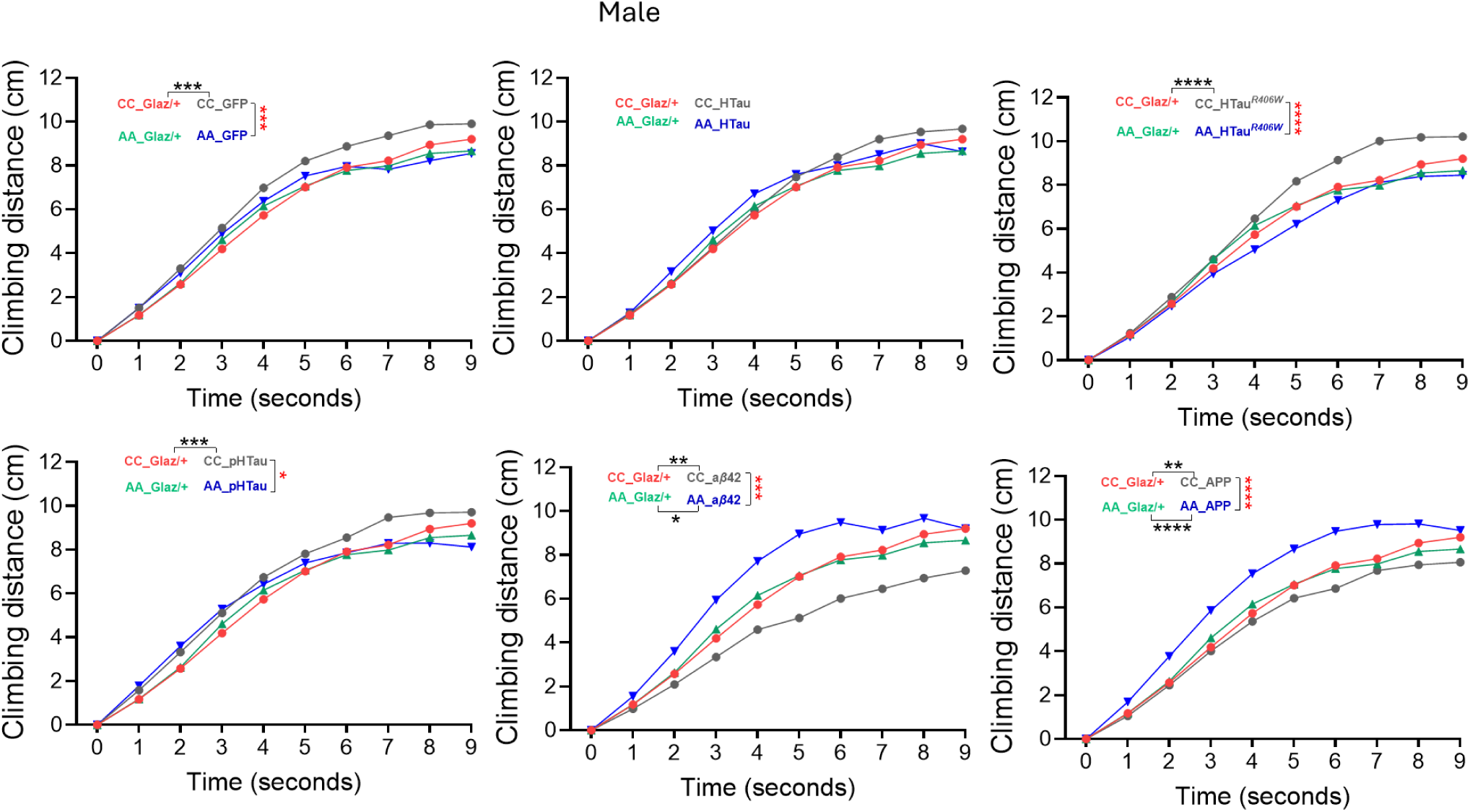
Male fly geotaxis performance during CC and AA condition from 0 to 9 seconds. Climbing distance showed no significant difference between AA and CC control flies. Whereas in AA flies GFP and Glaz/+ showed reduced (p<0.0001, p<0.001) compared CC GFP. HTau found no significant difference. R406W in CC showed a significant increase (p<0.0001, p<0.0001) compared to Glaz/+ and AA R406W. pHTau in CC showed increased compared to AA and Glaz/+ (p<0.03, p<0.001). AB42 in AA showed a significant increase (p<0.001, p<0.008) compared to Ab42 and Glaz/+ in CC. We also seen significant difference between AA Glaz/+ and AA AB42 (p<0.04). In APP, AA flies showed a significant increase (p<0.0001, p<0.0001) compared to AA, Glaz/+ and CC APP, in addition significant difference observed between CC Glaz/+ and CC APP (p<0.007). Two-way ANOVA followed by Sidak’s multiple comparisons test used for statistical analysis. n=3, p-values * <0.05, ** <0.01, *** < 0.001, **** <0.0001.

**SI. Figure 5.**
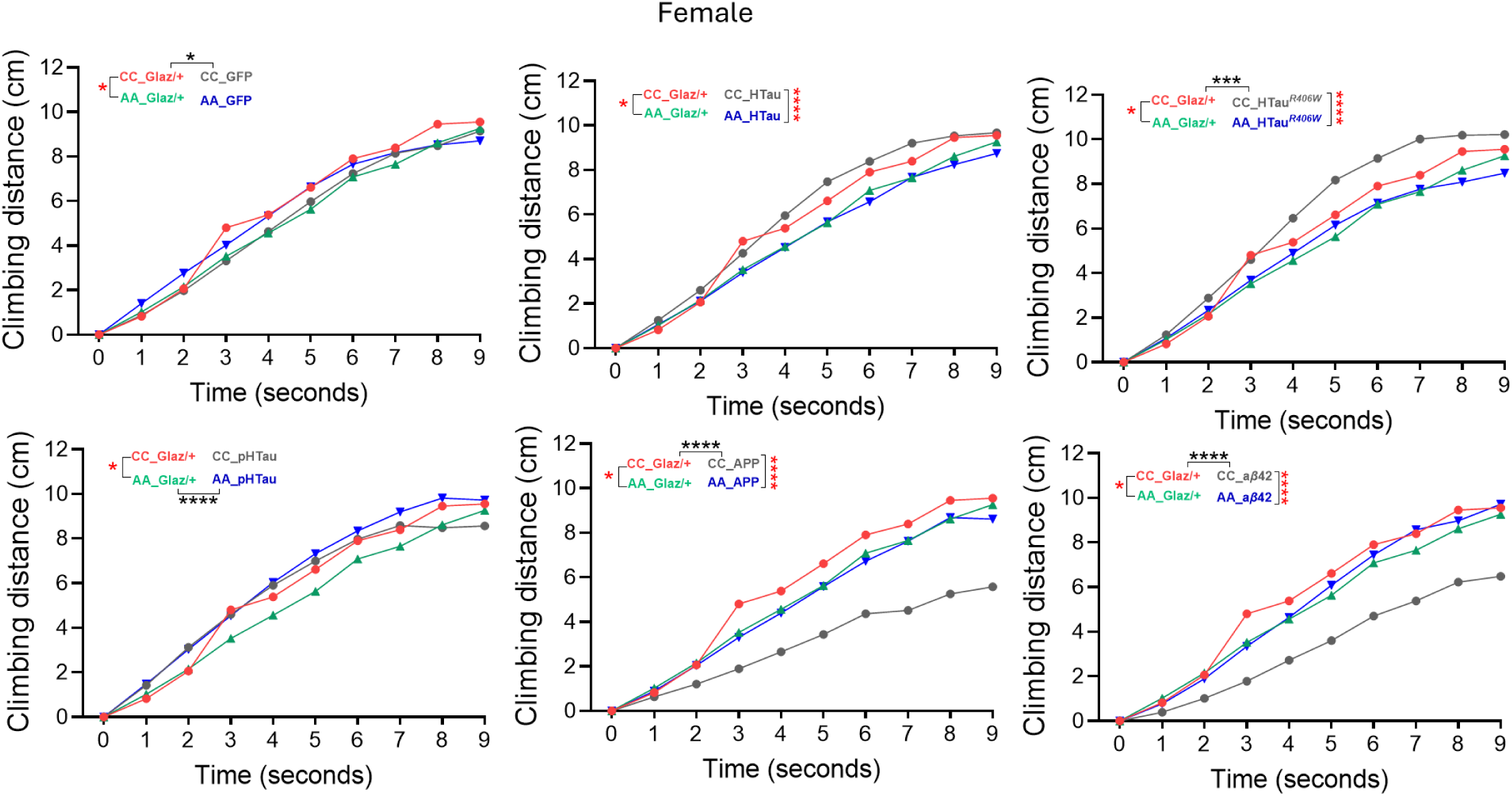
Female fly geotaxis performance during CC and AA condition from 0 to 9 seconds. Climbing distance showed significant difference (p<0.02) between AA and CC control (Glaz/+) flies. In CC condition Glaz/+ showed significant increase (p<0.04) compared to GFP. In HTau CC showed significant increase compared to AA (p<0.0001). R406W in CC showed a significant increase (p<0.001, p<0.0001) compared to Glaz/+ and AA R406W. pHTau in AA showed significant increase compared to AA Glaz/+ (p<0.0001). AB42 in CC showed a significant reduction (p<0.0001, p<0.0001) compared to Glaz/+ and AA AB42. APP in CC showed a significant reduction (p<0.0001, p<0.0001) compared to Glaz/+ and AA APP flies. Two-way ANOVA followed by Sidak’s multiple comparisons test used for statistical analysis. n=3, p-values * <0.05, ** <0.01, *** < 0.001, **** <0.0001.

**SI. Figure 6.**
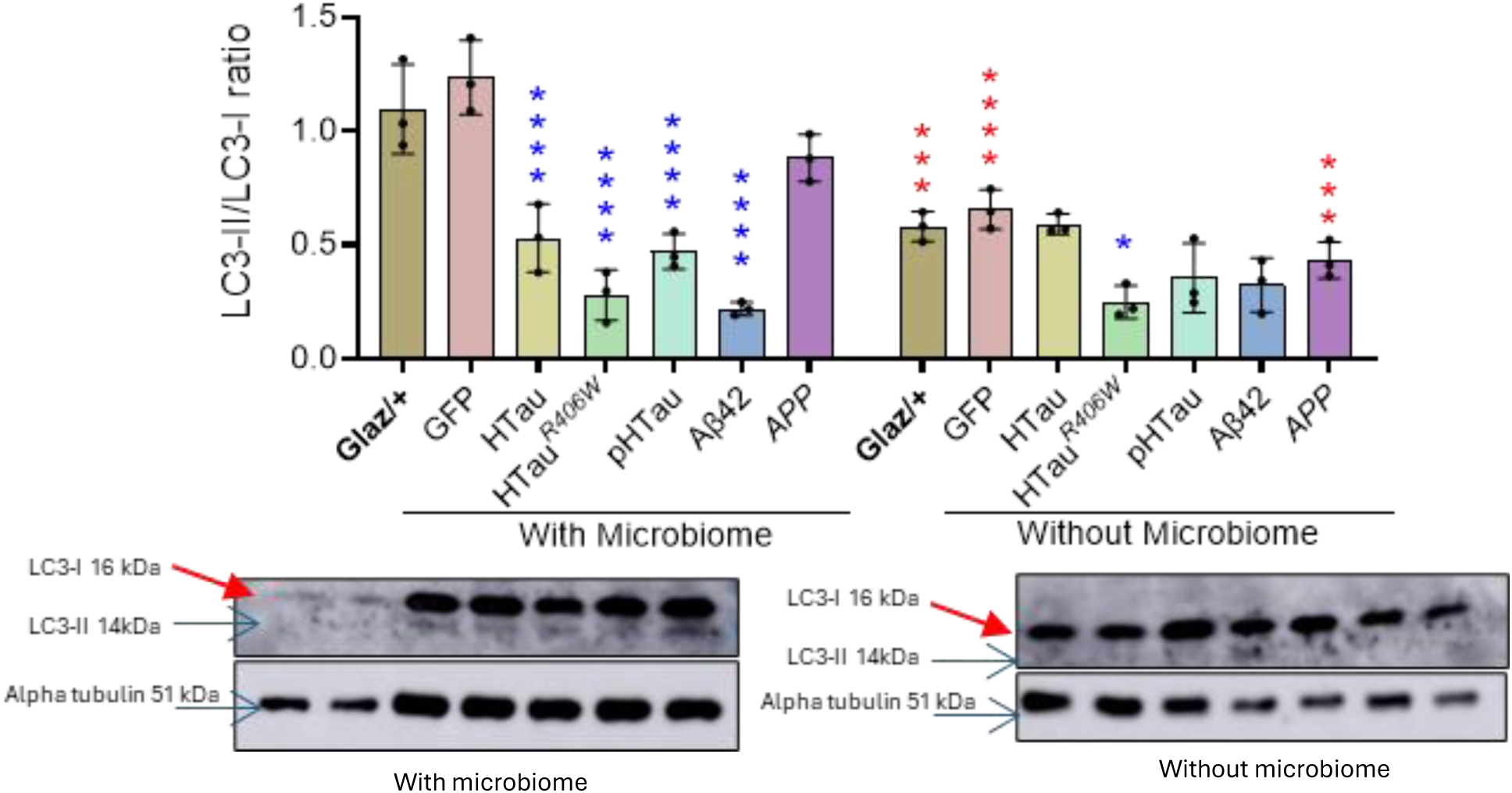
Representing the quantification data of LC3-II vs LC3-I ratio in transgenic flies. We compared the ration between two groups (with and without microbiome), also with in the groups we compared control vs transgenes. Our study showed a significant decrease in LC3-II expression in AD transgenes (with microbiome) compared to control and found a drastic reduction in axenic (without microbiome) condition. LC3 band intensity was normalized with alpha tubulin to understand the relative abundance of protein loading into the gels. Then we calculated the LC3-II/LC3-I ratio between genotypes and groups. Compared to Glaz/+, HTau, R406W, pHTau and AB42 showed a significant decrease (p<0.0001) in CC and R406W in AA (p<0.03) in each group. In AA flies Glaz/+ (p<0.001), GFP (p<0.0001), APP (p<0.001) showed a significant decrease compared to CC condition. n=3, p-values * <0.05, ** <0.01, *** < 0.001, **** <0.0001.

**Supplementary figure 7:**
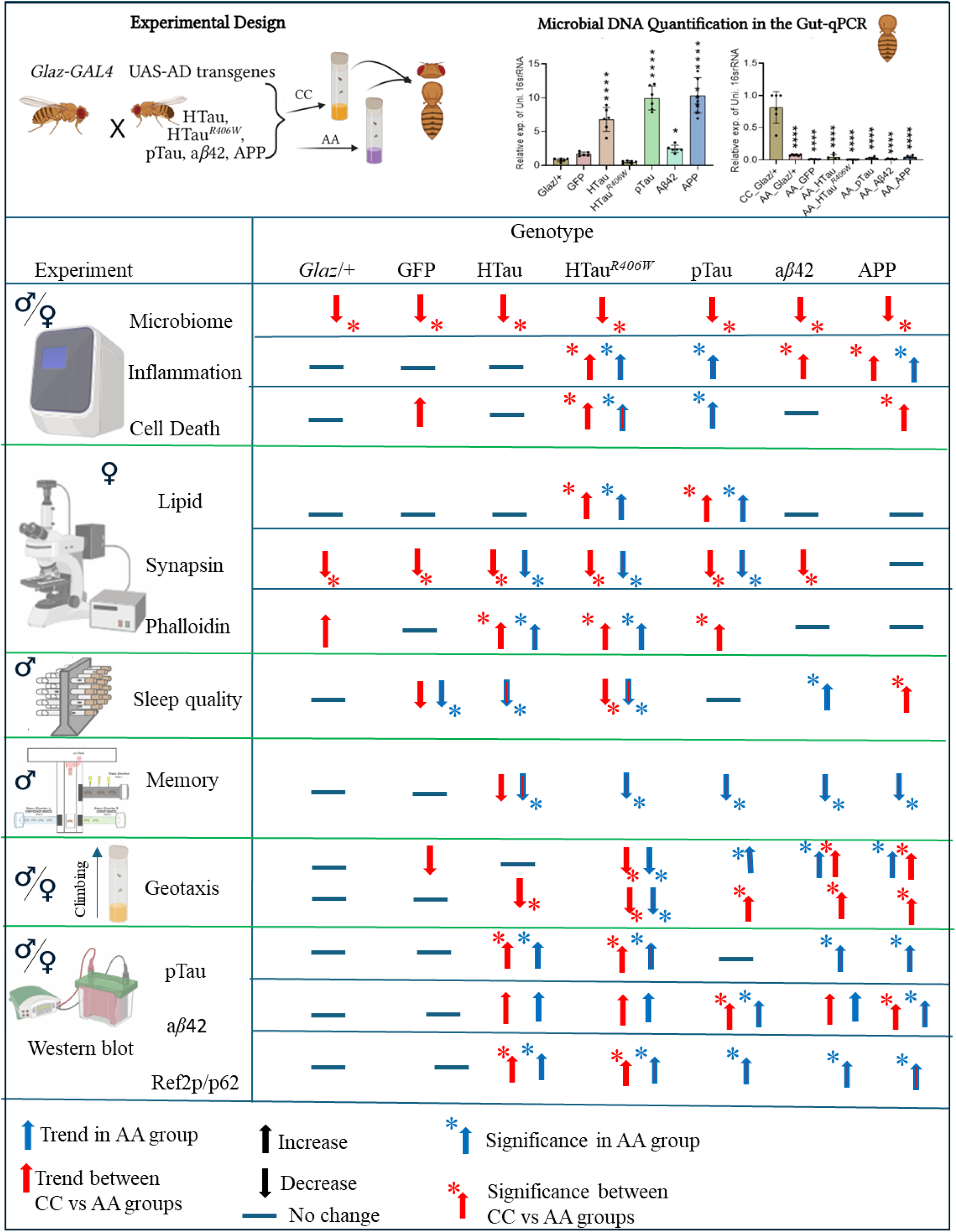
Summary Figure: Schematic representation of Work plan, and gut microbiome quantification using CC, AA flies. Experimental procedures including real time PCR for inflammatory, cell death markers and microbiome quantification. immunofluorescence analysis for lipid, synapsin and phalloidin staining. Sleep activity analysis with *Drosophila* Activity monitoring System (DAMS). Memory/Olfaction assay with T-maze. Geotaxis analysis and western blot analysis. Figure represents axenic fly data, compared with control (blue arrow) and between groups (CC vs AA, red arrow). Star represents significance. Up-arrow (Significant increase) Down-arrow (Significant decrease).

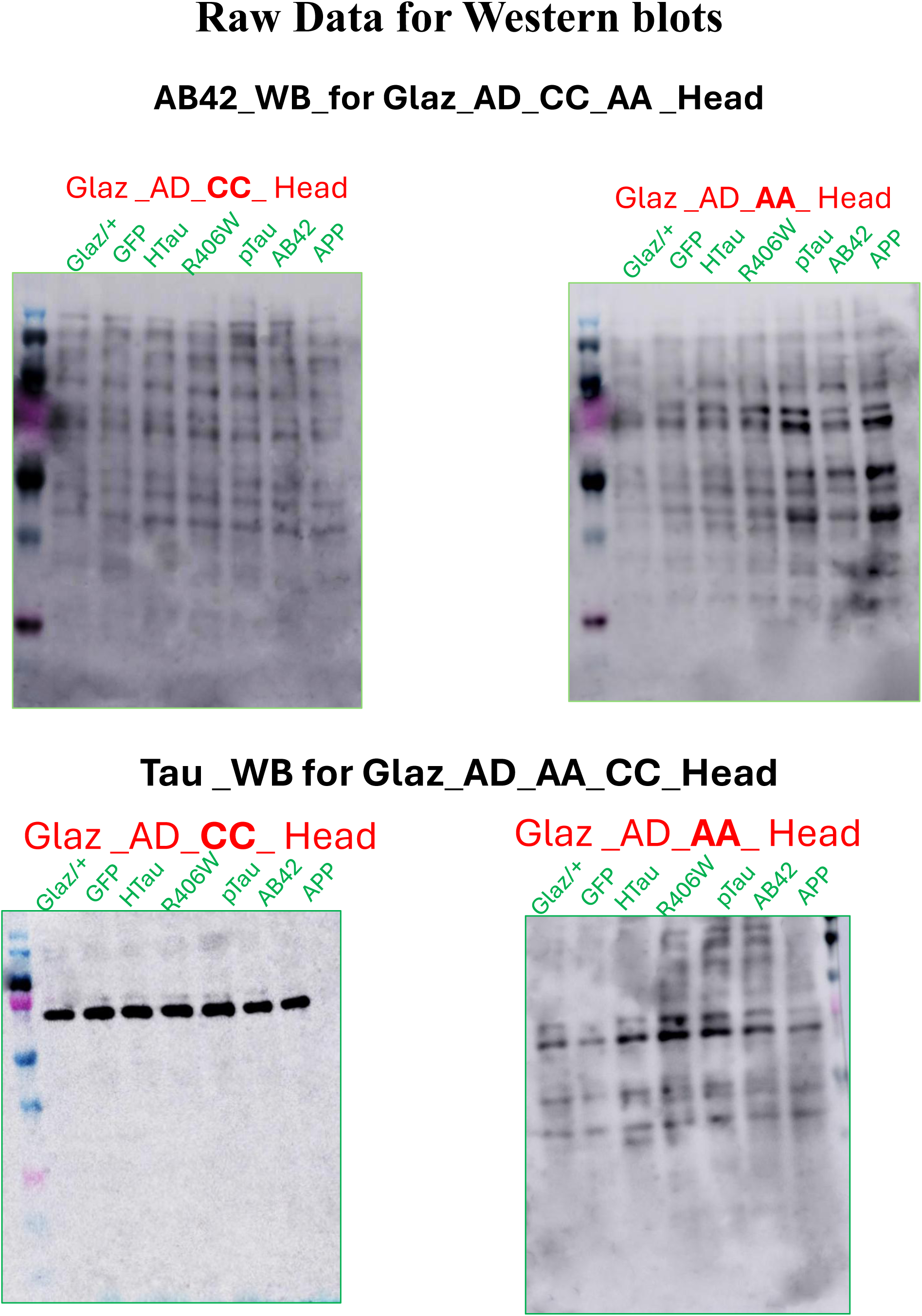

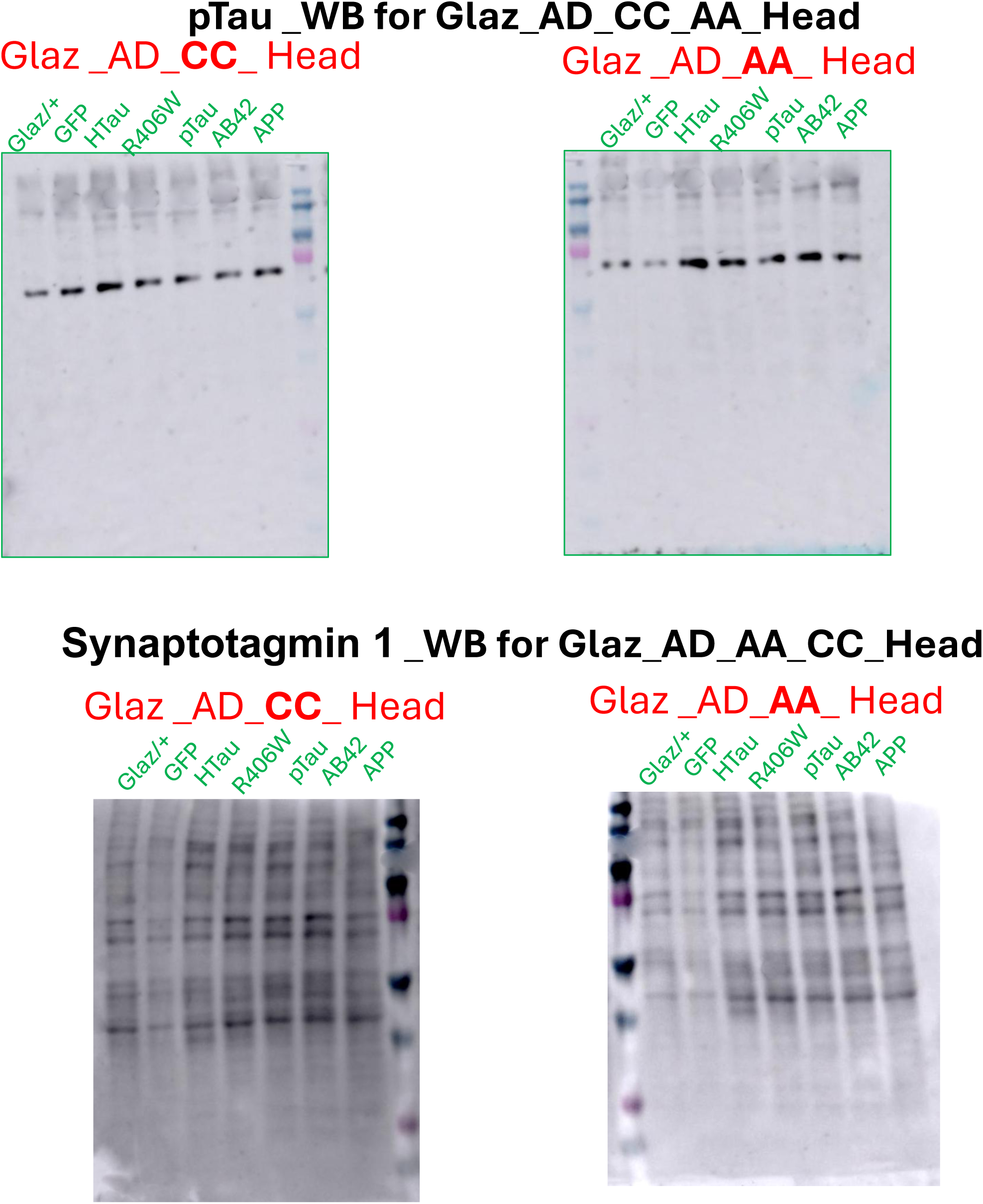

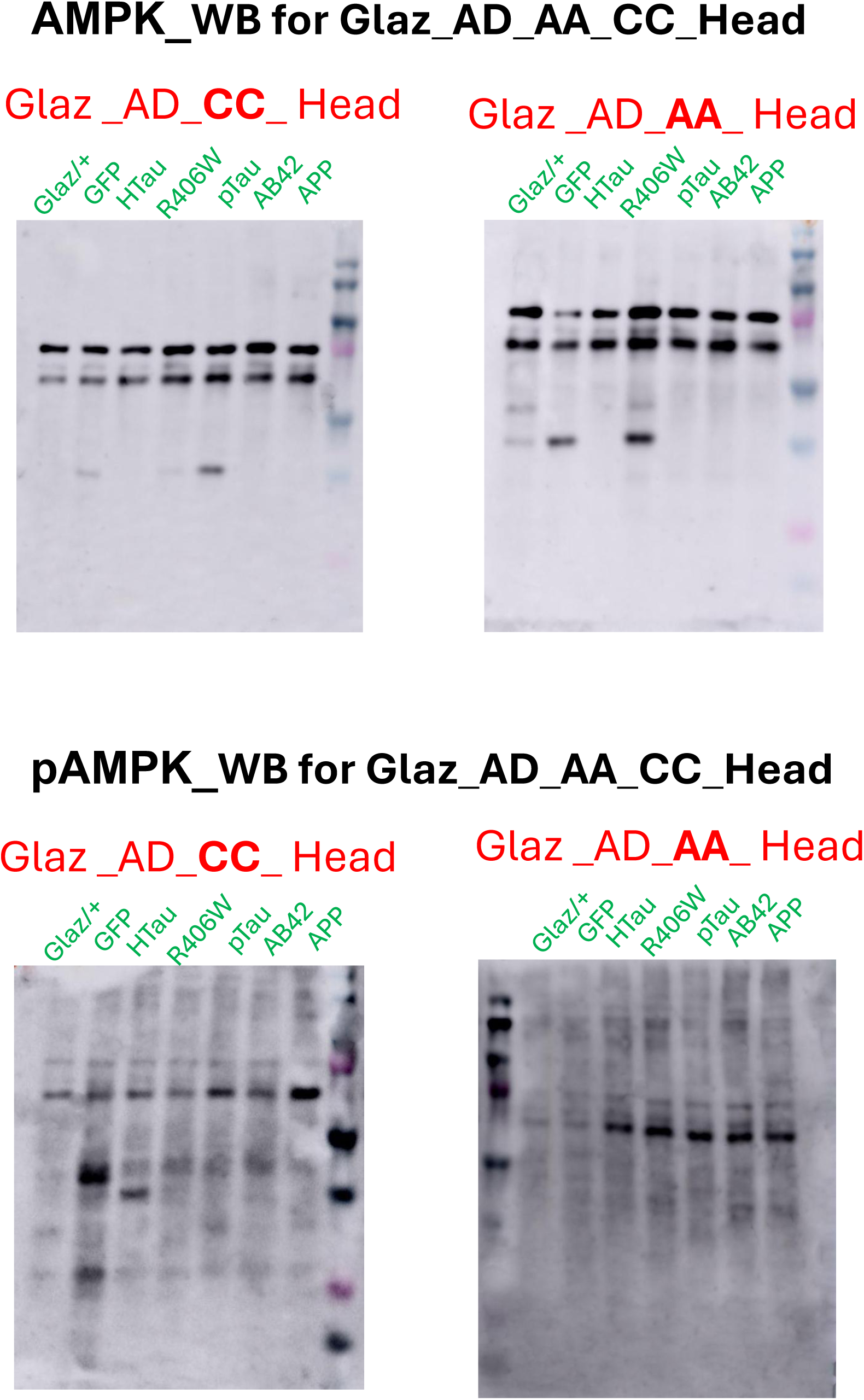

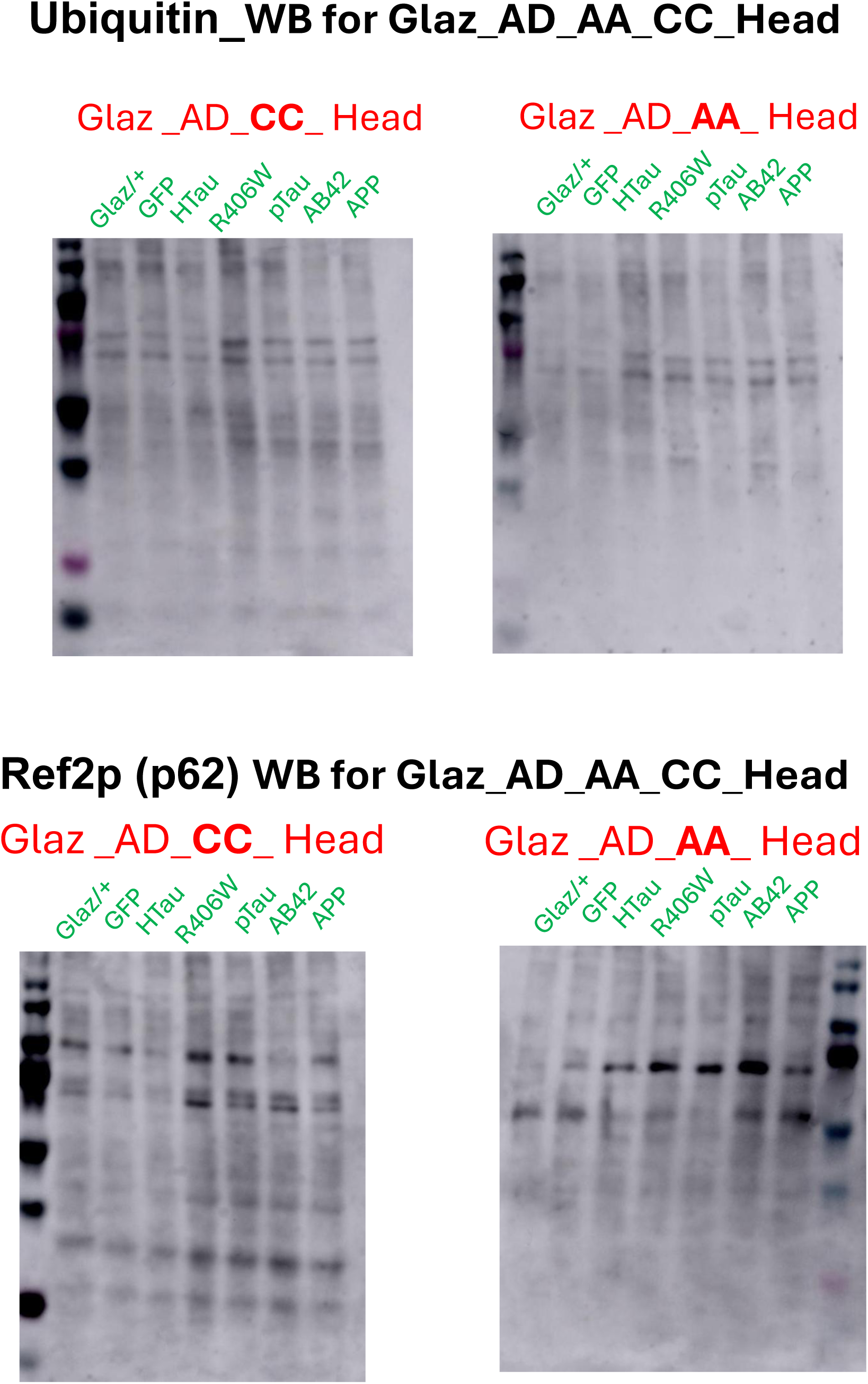

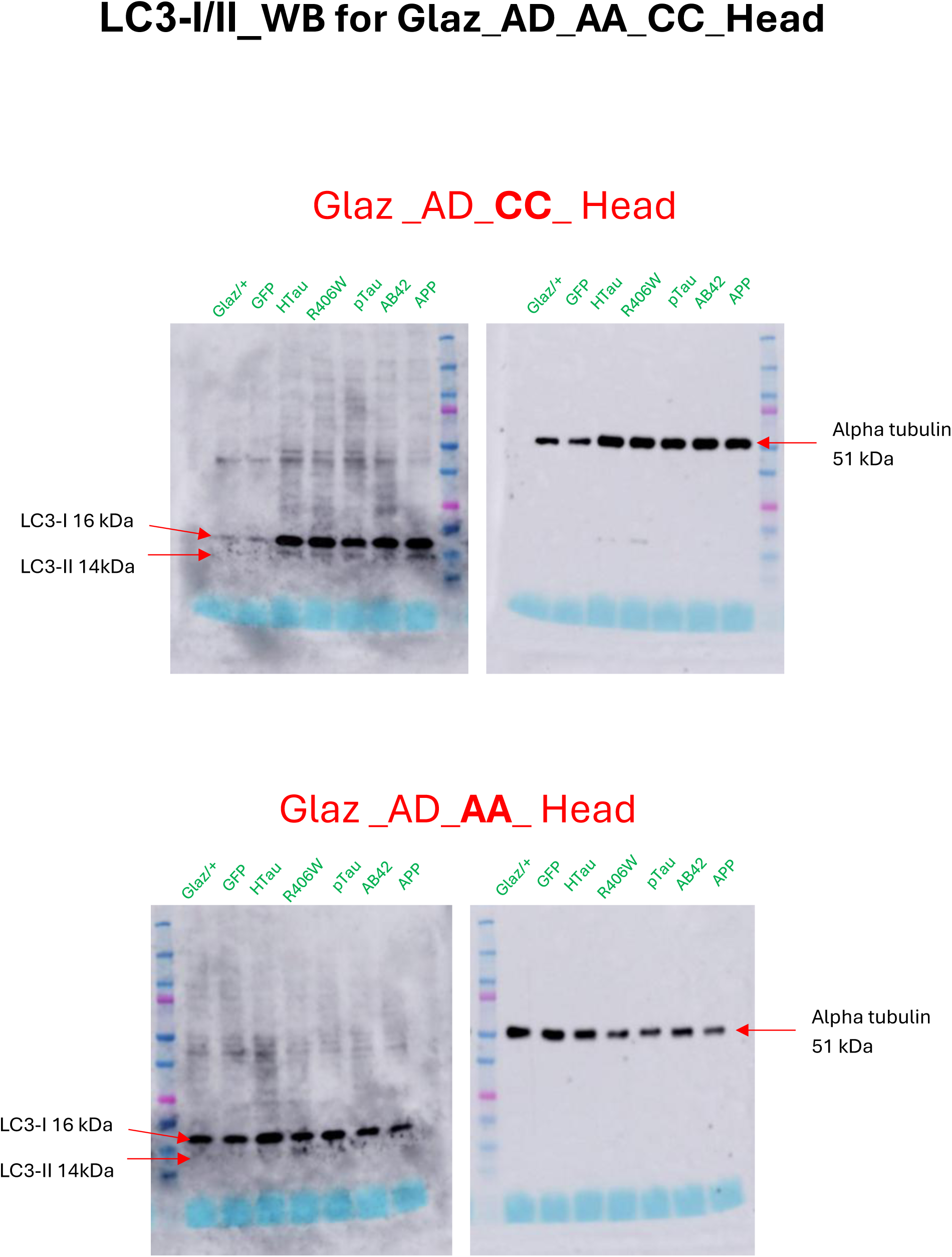

